# Defining specific cell states of MPTP-induced Parkinson’s disease by single-nucleus RNA sequencing

**DOI:** 10.1101/2022.05.29.493938

**Authors:** Yunxia Guo, Junjie Ma, Hao Huang, Jitao Xu, Kaiqiang Ye, Ning Chang, Qinyu Ge, Guangzhong Wang, Xiangwei Zhao

## Abstract

Parkinson’s disease (PD), a neurodegenerative disease with the impairment of movement execution that is related to age, genetic and environmental factors. 1-methyl-4-phenyl-1,2,3,6-tetrahydropyri-dine (MPTP) is a neurotoxin widely used to induce PD models, but the effect of MPTP on cell-gene of PD has not been fully elucidated. By single-nucleus RNA sequencing, we uncovered the PD-specific cells and revealed remarkable changes in their cellular states, including astrocytosis, endothelial cells absence, as well as a cluster of PD-exclusive medium spiny neuron cells. Furthermore, trajectory analysis of astrocyte and endothelial cells populations predicted candidate target gene sets that might be associated with PD. Notably, the detailed regulatory roles of astrocyte-specific transcription factors Dbx2 and Sox13 in PD were first revealed in our work. Finally, we characterized the cell-cell communications of PD-specific cells and found that the overall communication strength was enhanced in PD compared with matched control, especially the signaling pathways of NRXN and NEGR. Our work provides comprehensive overview on the changes of cellular states of the MPTP-induced mouse brain.

## Introduction

Parkinson’s disease (PD), a prevalent neurodegenerative disease, is predominantly characterized by motor disorders, followed by non-motor symptoms including cognition impairments, autonomic dysfunction and hyposmia^1^. PD mainly affects the elderly, accounting for a prevalence of 1.7% in population aged over 65 and the number of PD patients increases with aging, which causes serious health problems and care costs for the elderly and their families. Currently, PD is universally acknowledged to be caused by neuronal death in substantia nigra^2^, degeneration of dopaminergic neurotransmission, and the accumulation of a-synuclein (Lewy bodies) in neuronal cells^3^. However, PD is presently incurable, and the underlying mechanisms behind the neurological degeneration have been the subject of intense study over the last two hundred years.

Single cell/nucleus RNA sequencing (sc/snRNA-seq) technology has emerged as the most powerful instrumental for assessing cell type heterogeneity^4^, and this technique has been widely used in neuroscience. Currently, the majority of previous sc/snRNA-seq studies on PD were focused on iPSC-derived dopamine neurons^5, 6^ and mutant mouse (LRRK2, SNCA) postmortems brain^7–9^. However, most PD cases are sporadic, and up to 15% of PD cases are related to genetic mutations, but various environmental factors can also induce PD-like symptoms, including the exposure to pesticides or biotoxins. MPTP is a neurotoxin that can cause PD symptoms such as bradykinesia, postural instability, rigidity, cognitive deficits, and temporary autonomic disturbances^10^. MPTP can cross the blood-brain barrier (BBB) and be oxidized to 1-methyl-4-phenylpyridinium (MPP^+^) by monoamine oxidase B, and then MPP^+^ is concentrated in the dopaminergic terminals and cell bodies by the dopamine uptake transporter to produce toxicity^11^. This process is often accompanied by astrogliosis and microgliosis and endothelial cell injury^12^. However, all of these results were derived from traditional techniques such as immunohistochemical and positron-emission tomogram imaging^12^. In addition, current study on the MPTP-PD transcriptome is limited to RNA-seq for bulk tissues^13^. Although these studies provide valuable insights into the cellular phenotypic effects of MPTP on mouse brain, how MPTP affects the cell states at the single cell transcriptional level has yet to be elucidated.

Here, we applied snRNA-seq to investigate complex cellular state changes in the brain tissue of MPTP-PD and matched control (CN) mice. Firstly, we identified PD-specific astrocytes and endothelial cells based on cell proportion, cell density, differential expression genes and transcriptional regulation analysis. Then, the activation states of PD-specific cells were characterized by trajectory reconstruction analysis, and the gene sets that may mediate PD development were discovered. Moreover, another PD-related cells, PD-exclusive D2-medium spiniform neuron (D2-MSN), was identified through re-clustering of PD-deficient neurons, which might be an independent cellular state caused by MPTP induction. Eventually, we analyzed the the changes of communication relationship between PD-specific cells to explore the effects of MPTP on the communication pattern of these cells. Altogether, our work lays the foundation for elucidating the effect of MPTP on the cellular heterogeneity of brain tissue in PD, and we expect that our study will significantly facilitate future studies in PD mechanisms.

## Results

### Single-nucleus transcriptome profiling to identify cell populations

To investigate the effects of MPTP on the cellular heterogeneity of brain in PD, we performed snRNA-seq on the mixed samples of cerebral cortex, hippocampus, striatum and cerebellum from MPTP-PD and CN mice (Fig. S1a), which have been shown to be associated with PD in our previous work^13^. After filtering out potential doublets, poorly sequenced and damaged nuclei, 19,531 high-quality nuclei were kept for downstream integrated analysis (Fig. S1b, c). After batch correction, 24 clusters were identified and showed largely similar cellular landscapes in MPTP-PD and CN (Fig. 1a, Fig. S1d). By interrogating the expression patterns of known marker genes, eight major cell types were annotated: excitatory neurons (Ex1-13), inhibitory neurons (Inh1-4), astrocyte (AST1-2), microglia (MG), oligodendrocyte cell (OLG), oligodendrocyte precursor cell (OPC), endothelial cell (ENDO) and pericyte (PEC) (Fig. 1b, Fig. S2b and SM 1).

**Fig. 1.**
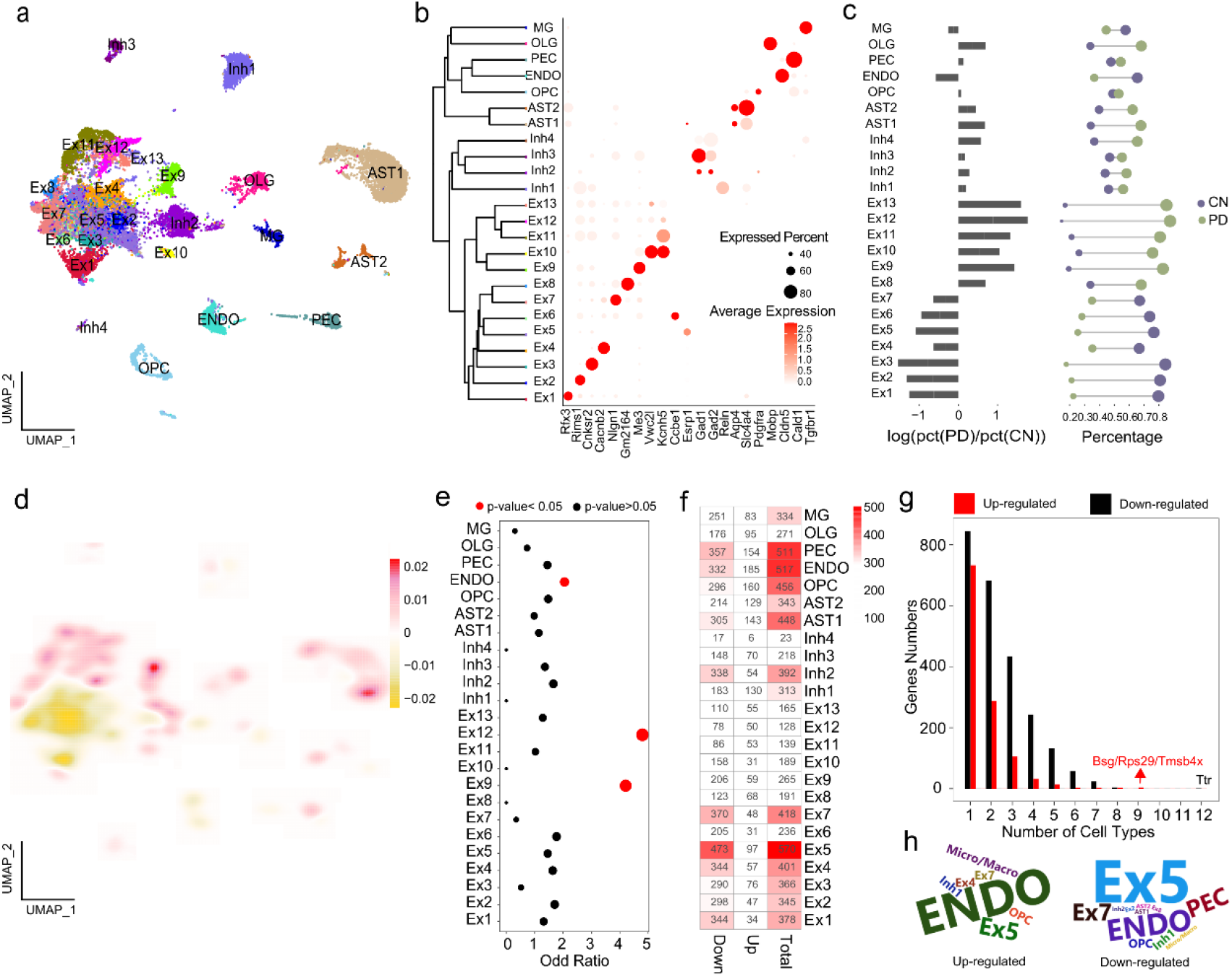
The characterization of cellular diversity in MPTP-PD brain by snRNA-seq. **a,** Contribution of nuclei from PD or CN to each cell type; colored by cluster. **b,** Cell representative marker genes. Expression level (color scale) of marker genes across clusters and the percentage of cell expression (dot size). **c,** The changes in frequency of multiple cell types between PD and CN. Left: log ratio of average fraction in PD vs CN. Right: proportion of PD and CN profiled cells. The color and dot size represent different samples and the percentage of cells, separately. **d**, Differential 2D cell density between PD and CN. Red and yellow indicate the high and low density of cells in PD, respectively. **e,** Using Fisher’s exact test to get cell types in which PD-risk gene enrichment. Circle size indicate OR value, and red color highlight enriched cell types with p-value <0.05. **g,** The number of up- or down-regulated genes per cell type was detected. **h,** PD-risk DEGs detected frequency in each cell type.

The results of cell correlation further confirmed the accuracy of cell classification (Fig. S2a). To investigate the changes in cell-type composition associated with MPTP-PD, three approaches were used. Initially, we examined changes in the composition of each cluster in the context of disease and found several that were overrepresented (Ex8-13, AST, OLG) or underrepresented (Ex1-7, ENDO) in MPTP-PD, and the proportion of other cells was similar to CN (Fig. 1c). Subsequently, we compared PD and CN cell density distributions in the UMAP representation, and found that the fraction of AST1, Ex4 and Ex9 in MPTP-PD were increased compared to CN (Fig. 1d, Fig. S2c). In addition, we also evaluated whether the expression patterns of PD-associated risk genes were cell-specific. The results showed that PD risk variants were significantly enriched in Ex9, Ex12 and ENDO cells (p-value < 0.05, OR > 1) (Fig. 1e). Altogether, these results preliminarily predicted that AST1, ENDO and excitatory neuron might be the cell types with the most obvious effects of MPTP on PD.

### Multi-dimensional validation of MPTP-PD specific cells

To verify our prediction of PD-specific cells, we further assisted from multi-dimensions including differential expression genes (DEGs) and transcriptional regulation analysis. We compared the numbers of cell-types DEGs between PD and CN, and found that about 75% of DEGs were down-regulated in PD, especially in Ex1-7, AST1, and ENDO (Fig. 1f). The detected frequency analysis showed that the DEGs had strong specificity in each cell types, and the number of detected down-regulated genes were more than that of up-regulated genes in PD (Fig. 1g). The down-regulated gene Ttr could be identified in the half of cell types, while up-regulated genes Bsg, Rps29 and Tmsb4x were detected in 9 clusters (Fig. 1g), and all of them were dysregulated in ENDO and Ex5 (SM 2). Of that, Cakar *et al.* revealed that polyneuropathy can be caused by the accumulation of amyloidogenic TTR protein in tissues, as in Alzheimer’s disease (AD) and PD^14^. P53 mediates cell defects associated with Rps29, and p53 inhibitors were very effective in maintaining motor function in PD mice^15^. Overexpression of Tmsb4x in cultured hippocampal neurons can reportedly reduce neurite outgrowth and neuronal development^16^. Notably, a greater number of PD risk DEGs were obtained in ENDO and Ex5 cells in MPTP-PD (Fig. 1h).

Transcription factors (TFs) tightly control cell fate in neurodevelopment and have been implicated in neurodegenerative processes^17^. Therefore, we validated MPTP-PD specific cell types from the perspective of transcriptional regulation, and further explored the effect of TFs on disease. We identified 213 and 293 significant TFs in MPTP-PD and CN respectively, and the most of the CN and PD specific TFs were contributed from Ex5 and AST1 (Fig. 2a). The heatmap of top3 specific TFs of each cell type showed the activation status of specific regulatory factors in each cell type, among which only 10 TFs (Bhlhe22, Lhx9, Ovol2, Cux2, Uncx, Rarb, Sox9, Emx2, Tbx2, Nr1h3) were co-activated in the same cell types of PD and CN, but the regulatory intensity was different (Fig. 2b). It suggested that alterations in the activation of TFs may drive changes in disease cell states. Subsequently, we focused on 155 overlapped TFs from all clusters between PD and CN and found that 101 (65%) co-regulated conserved TFs have significant similar activated states among all clusters between PD and CN, while the remaining TFs have strong cell-specific activated patterns in PD or CN (Fig. S3). It was suggested that the activation of some TFs with cell type specificity might be revealed the changes of intracellular transcriptional regulatory network, thus affecting the development of PD. Finally, the activation status of rest 54 TFs suggested that there was a general homogeneity in the activation or inhibition of TFs in all cell types, only Rarb and Foxp2 were simultaneous activated and inhibited in different cell types, and Maf and Xbp1 were activated or inhibited in almost all neurons of PD, respectively (Fig. S4). In conclusion, we systematically revealed candidate *trans-*regulatory elements in different cell types of MPTP-PD for the first time, especially disease-related AST1.

**Fig. 2.**
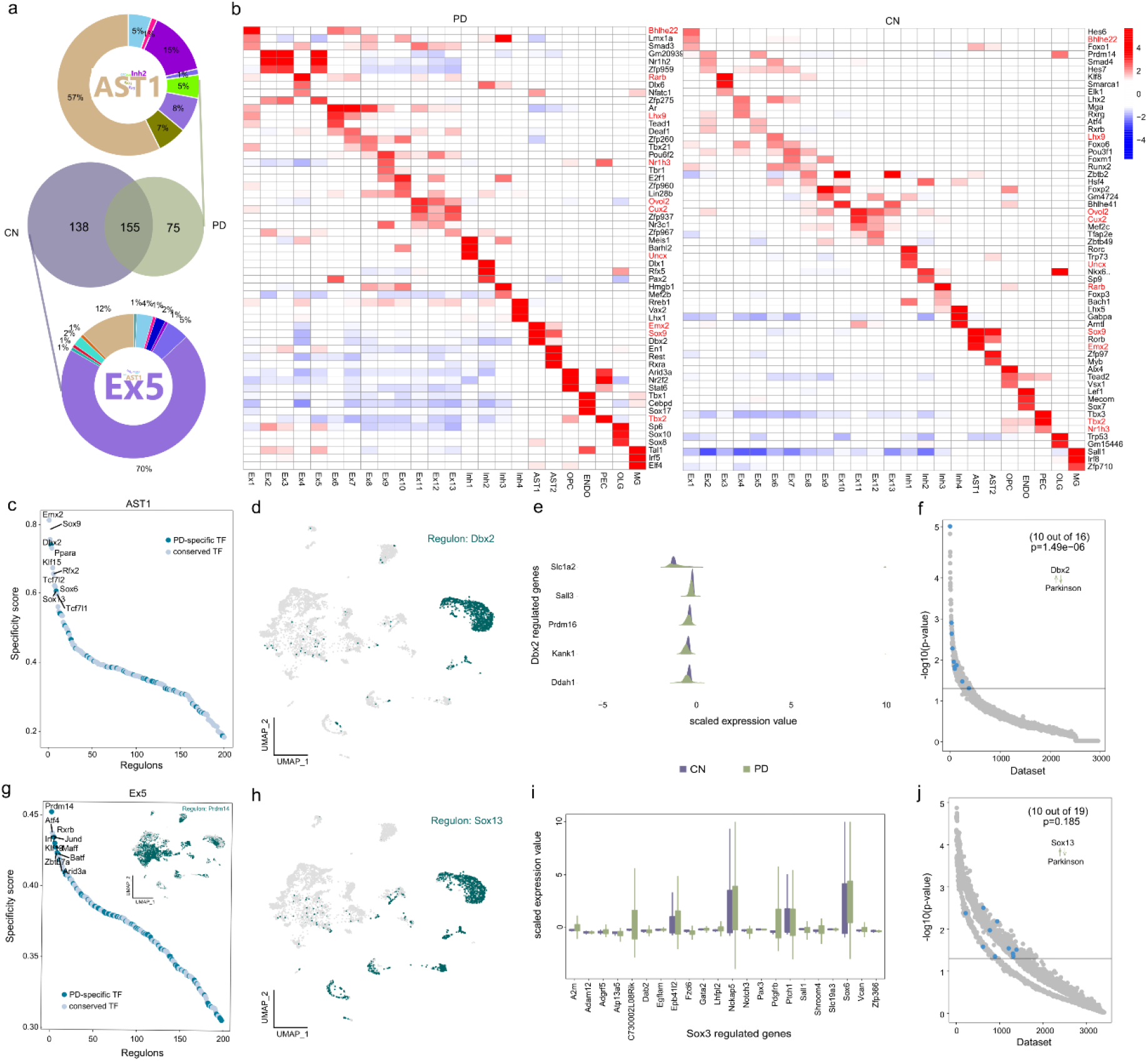
Cell type-specific transcription factors in disease. **a,** Venn plots of conservative and specific TFs in PD and CN. Middle venn plot shows conservative and specific TFs detected in PD and CN. The circle percentage diagram with word cloud insert detailed present specific TFs of CN and PD were mainly derived from Ex5 and AST1, the colors of circle diagram and word cloud plot correspond. **b,** Heatmap of top three specific TFs for each cell types in PD (left) and CN (right). **c, g,** The rank of regulons in AST1 and Ex5 based on regulon specificity score. **d, h,** Binarized regulon activity scores for top regulons Dbx2 and Sox13 on UMAP map (dark green dots). **e, i,** Expression levels of Dbx2 and Sox3 transcription regulated target genes in AST1. **f, j,** SEEK coexpression result for target genes of top regulons Dbx2 and Sox13 in different public datasets. The x axis represents different datasets, and the y axis represents the co-expression significance of target genes in each dataset, AST1 related datasets with significant correlation (p-value < 0.01) are highlighted by blue dots.

Taken together, the results of cell proportion, cell density and PD-risk gene enrichment analysis preliminarily predict that AST1, ENDO cells and neuron cells might be associated with PD. Subsequent results of DEGs and transcriptional regulation further verified our hypothesis. For example, the number of down-regulated genes in EX1-7, AST1 and ENDO cells was prominent, the up- and down-regulated PD-risk genes were mainly from Ex5 and ENDO cells, and 57% of PD-specific TFs were from AST1. Therefore, our multi-dimensional methods ultimately focused on PD-specific cell types: AST1, ENDO and PD-deficient neurons, which will be the focus of further research.

### Transcriptional regulation of disease-specific astrocytes

To investigate the specific TFs of AST1 and Ex5 cells and their regulatory roles in PD, we sought to evaluate the cell-specific TFs activation states. Dbx2 and Sox13 were identified as the most prominent specific TFs associated with AST1 in MPTP-PD (Fig. 2c), and Prdm14 was the top TF that associated with Ex5 in CN based on the rank of regulon specificity sore (Fig. 2g). UMAP plot provided additional support that the activities of Dbx2 and Sox13 were highly specific to AST1 (Fig. 2d, h), but Prdm14 was not only activated in Ex5 (Inset of Fig. 2g). Subsequently, the genes regulated by Dbx2 and Sox13 were identified by RcisTarget^18^, and the expression levels of these genes were investigated in PD and CN, respectively. The genes that regulated by Dbx2 were under-expressed in PD, while the expression of Sox13 regulated genes were opposite, indicating that Dbx2 and Sox13 may act as transcriptional inhibitor and activator in MPTP-PD (Fig. 2e, i). To further evaluate the accuracy of our findings, we applied SEEK analysis to search for GEO datasets about the co-expression pattern of Dbx2 and Sox13 target genes, then highlighted the work tittle that co-occur with the term ‘Parkinson’, and found that these co-expressed genes tend to be associated with PD (Fig. 2f, j and SM 3). For example, Slc1a2, Prdm16, Kank1 and Ddah1 were target genes of Dbx2, the high expression of Slc1a2 was found to reduce the risk for PD in a Chinese cohort^19^, and the other genes are associated with cognitive function^20^, autism spectrum disorder^21^ and depression-like behavior^22^, respectively. Meanwhile, Sox13 target genes Ptch1, Pdgfrb, Vcan, Lhfpl2 and A2m have been reported to be related to PD. Other regulated genes by Sox13 might be associated to PD syndrome, such as Nckap5 is considered the most promising candidate for bipolar disorder^23^, and Epb41l2 gene is associated with cognitive impairment in the hippocampus induced by anesthesia^24^. So far, the study about Dbx2 and Sox13 has focused on neural stem cells^25^, and the role of Dbx2 and Sox13 in PD has not been studied.

### Trajectory reconstruction of MTPT-PD associated astrocytes and endothelial cells

To investigate the changes of AST1 and ENDO cells states in MPTP-induced mice, we subclustered these cells and reconstructed their activation trajectories. We identified five AST1 subpopulations characterized by high expression of Meg3, CT010467.1, Apoe, Lsamp, and Luzp2 (Fig. 3a, Fig. S5a). Subsequently, we reconstructed a cell trajectory structure comprising these major sub-populations using the DDRTree method of Monocle3^26^. The activation trajectory of AST1 spans from Meg3^High^ cells towards two activation branches, one containing Apoe^High^ cells and the other with clusters highly expressing Luzp2 and Lsamp (Fig. 3a). It has been reported that the relative expression level of Meg3 in PD patients is lower than that in healthy population^27^, while Apoe has an impact on cognitive decline of PD^28^. Luzp2 is found to be associated with AD^29^, schizophrenia^30^, intelligence^31^, and verbal memory^32^, and the level of Lsamp is increased both in patients with depression and schizophrenia^33^. The result highlight that these genes may be linked with PD or its syndrome. Importantly, we observed that these five subclusters were all distributed in AST1 cells of UMAP, especially the cluster with high Luzp2 and Lsamp expression were the cells of increased AST1 in cell density analysis (Fig. 1e, Fig. S5b). We observed that Luzp2 and Lsamp genes were distributed in the hippocampal based on the results obtained from in situ hybridization of Allen Brain Atlas (Fig. S5c). To further characterize the linked genes of these activated AST1 states in PD, we identified 100 genes whose expression was associated with the activation trajectory, of which 42 and 48 genes were independently highly expressed in CN and PD, respectively (Fig. 3b). To identify the potential pathways associated with marker genes of Lsamp^High^ and Luzp2^High^ subclusters, we performed functional annotation using hypergeometric test based on Kyoto Encyclopedia of Genes and Genomes (KEGG) database. These two subpopulations were associated with locomotory behavior, trans-synaptic signaling of endocannabinoid, response to auditory stimulus and glutamate receptor pathways (FDR < 0.05) (Fig. 3c). Next, we performed functional annotation for PD up- & down-regulated genes (Fig. 3d and SM 4) in all AST1 subpopulations based on the molecular function of Gene Ontology (GO) database. The results showed that PD-up-regulated genes were associated with synaptic, dendritic/neuron spine, startle response, synaptic transmission and ion channel regulator activity (FDR < 0.05) (Fig. S5d), which were associated with PD in previous studies^34–36^. Meanwhile, 14 overlapped genes were obtained between the DEGs and the activation-trajectory associated genes in AST1 (Fig. 3e and SM 4). Although none of these genes overlapped with existing PD-risk gene sets, most of them were candidate genes related to autism^37^, dyskinesia^38^, schizophrenia^39^. We speculated that these genes might be involved in the development of PD.

**Fig. 3.**
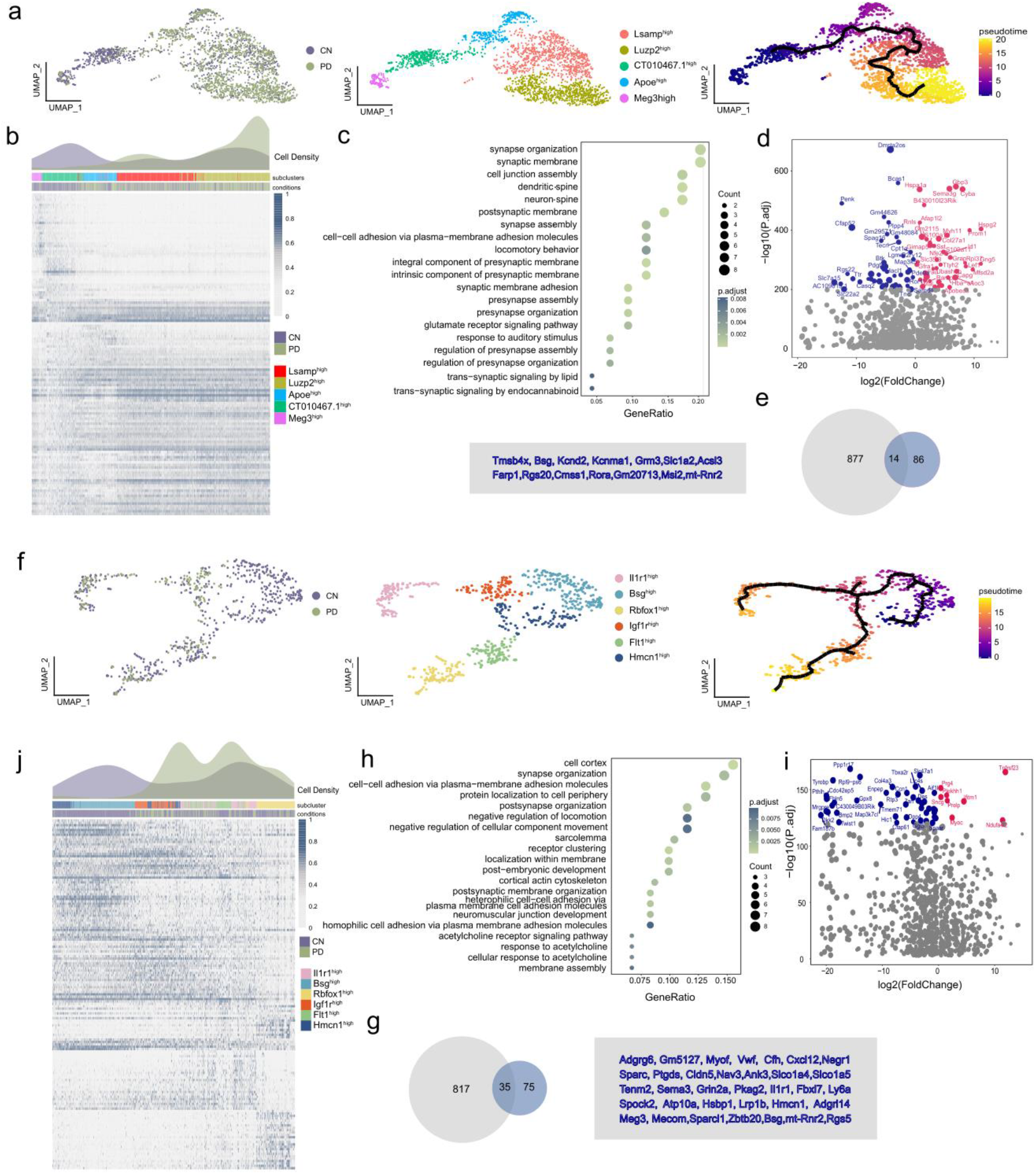
Trajectory reconstruction reveals astrocyte and endothelial differential activation in PD. **a, f,** AST1 and ENDO subclusters labeled with a representative marker gene and trajectory reconstruction and pseudotime representation of subclusters. **b, j,** PD and CN differential cell-density distribution along pseudotime. The expression of 100 and 113 genes highly associated with the AST1 and ENDO activation trajectory, respectively. **c, h,** KEGG and GO terms associated with genes of the Luzp2^High^ and Lsamp ^High^ cells in AST1 and Hmcn1^High^, Igf1r^High^, Flt1^High^ in ENDO, respectively. **d, i,** Volcano map of DEGs in PD and CN. The upregulated genes with red dots, downregulated genes with blue dots. **e, g,** The overlapped genes between PD-DEGs and the DEGs along the AST1 and ENDO activation trajectory

Following the same analytical approach mentioned above, we identified six ENDO subclusters characterized by high expression of Hmcn1, Bsg, lgf1r, ll1r1, Flt1 and Rbfox1 (Fig. 3f, Fig. S6a, b), and recovered their activation trajectory. The results implied an ENDO activation transited from Hmcn1^High^ to Rbfox1^High^ and ll1r1^High^ subclusters (Fig. 3f). We observed that ENDO cells were generally absent in PD, but the cells of Bsg^High^ was most severely, which was almost completely deletion (Fig. S6c). The Bsg gene is specifically expressed in ENDO of brain, and it has been reported that Bsg knockout mice exhibited deficits in learning and memory^40^. Indeed, ENDO cells in PD were highly enriched at the two activation branches of their trajectory (Rbfox1^High^ and ll1r1^High^) compared to CN (Fig. 3j, Fig. S6d). Rbfox1 is one of the risk genes that are common to PD and various psychiatric disorders^41^. ll1r1 can regulated by miRNAs that have been implicated in the potential regulators of alcohol-related neuroinflammation, inducing brain injury and neurodegeneration^42^. Subsequently, we performed functional annotation for Hmcn1^High^, Igf1r^High^ and Flt1^High^ subcluster marker genes based on the GO database, the results showed that they were highly functionally related to negative regulation of locomotion and cellular component movement, cell adhesion and acetylcholine receptor (FDR < 0.05) (Fig. 3h). The acetylcholine receptors may be stimulated by endogenous agonists such as acetylcholine, or exogenous chemicals such as nicotine, to activate physiologic angiogenesis or pathologic angiogenesis^43^. Next, we identified 35 overlapped genes between DEGs and activation locus genes of ENDO cells (Fig. 4i, g and SM 4). About half of the overlapped genes have been found to be related to PD in previous studies, and the rest genes are related to neuropsychiatric diseases (Zbtb20, Nav3), tissue aging (Myof), impaired memory (Atp10a), blood-brain barrier (Slco1a4/5, Cldn5, Ly6a)^44–46^.

**Fig. 4.**
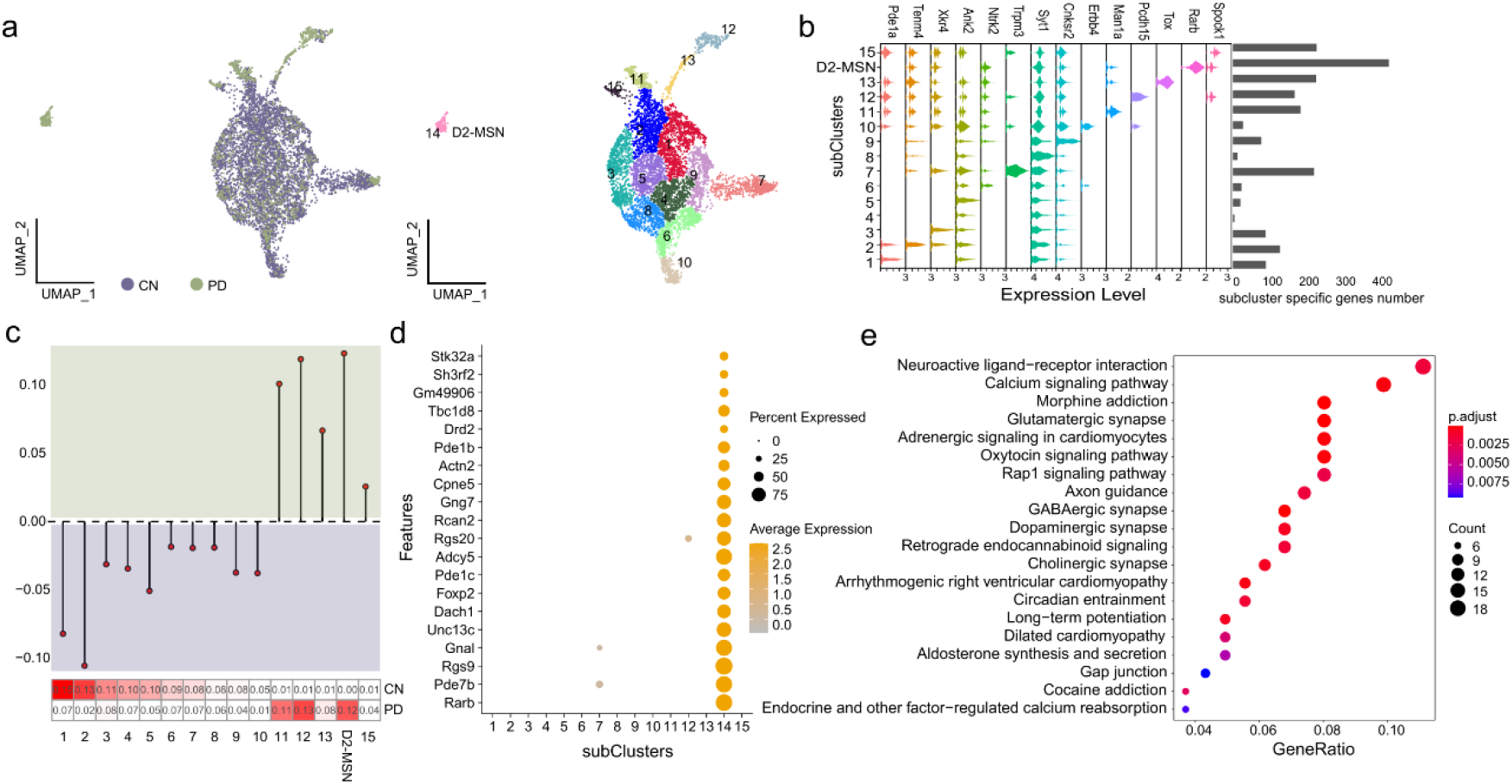
Neuronal states specific to disease models. **a,** UMAP dimensionality reduction of Ex1-8 from the snRNA-seq analysis. **b,** Marker gene expression and the number distribution of cluster-specific higher expressed genes. **c,** The proportion of CN and PD in 15 subclusters. Top: the lollipop of CN vs PD, y>0 means that the proportion of cells in PD is greater than CN; Bottom: heatmap of the cells proportion in PD and CN. **d,** bubble chart of all marker gene expression in subcluster 14. **e,** KEGG terms associated with genes of sbucluster14.

Since previous study indicated that OLG and OPC were related to PD^9^, we also performed trajectory analysis for OLG and OPC cells. Compare with AST1 and ENDO, the trajectory and cell density distributions analysis of OLG and OPC showed more similar states, and the heatmap of trajectory-dependent genes density also showed the similar pattern. (Fig. S7).

### Characterization of transcriptomic state of neuron cells

In order to explore the effect of MPTP on neurons in the process of inducing PD, we further deciphered the identity of the neuron, which were absent in PD. We separated all excitatory neuron cells into two major groups, Ex1-8 and Ex9-13, according to gene expression similarity (Fig. 1c Hierarchical cluster diagram). Almost all subclusters in Ex1-8 showed absence of PD cell except for Ex8 (Fig. 1d). We re-clustered Ex1-8 to distinguish 15 subclusters (Fig. 4a), and surprisedly found that subcluster14 consisting of 206 cells had no continuity with other subclusters and only concentrated in PD (Fig. 4a, c). Moreover, subcluster14 was derived from cells with increased Ex4 in cell density distribution analysis (Fig. S8a). The number distribution of cluster-specific higher expressed genes showed that subcluster14 had the most marker genes (Fig. 4b). And the top four highly expressed genes of subcluster14 could be verified in the striatum of Allen Brain Atlas (Fig. S8b). Among them, the mutations in Rarb could cause intellectual disability with progressive motor impairment^47^, and Pde7b plays an important role in schizophrenia^48^ and dopaminergic cell death^49^. Rgs9 is a potent modulator of G-protein-coupled receptor function in striatum^50^, and dopamine receptors are associated with distinct G-proteins^51^. The mutations in the Gnal could cause primary torsion dystonia^52^. In addition, some other marker genes of subcluster14 have been extensively studied. The recent literature report indicated Dach1 expression in the human striatum MSN^53^.

Adcy5 mutations have been associated with substantia nigra damage, and white and gray matter changes in striatal cortical pathways^54^. Expression levels of Rcn2 was responsive to external stressors such as reactive oxygen species, Ca^2+^, amyloid beta, and hormonal changes and upregulated in degenerative neuropathy^55^. Gng7 is the abnormal protein of dopaminergic signaling^56^. CPne5 is the circadian rhythm-related proteins, and circadian rhythm have a direct or indirect effect on the neurodegenerative processes^57^, and more importantly, the gene is involved in PD-induced toxins like paraquat^58^. In accordance with the expression characteristics of these genes (Fig. 4d), we defined subcluster14 as the PD-exclusive D2-MSN that was located in the striatum. Finally, we performed KEGG and GO analysis on the marker genes of D2-MSN, and found that all terms are significantly associated with neuronal synapse (e.g., dopaminergic, glutamatergic, cholinergic, GABAergic), ligand-receptor interaction pathway, ion channel and other related functions (FDR < 0.05) (Fig. 4e, Fig. S8c). For example, study has shown that a-synuclein can induce dysregulation of miRNAs, which target the neuroactive ligand-receptor interaction pathway^59^.In conclusion, combined with functional analysis and literature review of marker genes obtained from subclusters14, we speculated that this subcluster was in an independent cell state during MPTP induction.

### Analysis of cell-cell communication in MPTP-PD specific cells

Integrating pathways and functions of all PD-specific cells speculated that MPTP was likely to alter cell-cell communication. For example, the top enriched terms in D2-MSN included neuroactive ligand-receptor interaction and calcium signaling pathway, synaptic membrane and cell junction assembly in AST1, cell-cell junction in ENDO and so on. To further explore the interactions between PD-specific cells, we applied CellChat to infer intercellular communication networks. The changes of cell communication analysis require the same cell population composition between two datasets. Thus, we first used AST1 and ENDO cells between PD and CN. We found that the global number of ligand-receptor (L-R) pairs was decreased in PD, while the interaction strength was enhanced in PD compared to CN (Fig. 5a, b). Interestingly, although ENDO_Rbfox1^High^ and AST1_Meg3^High^ cells were reduced in PD, the number and intensity of intercellular communication were most significant (Fig. 5b). Next, we were curious about which signaling pathways and ligand-receptor pairs (L-R pairs) changes cell communication network. We further compared the information flow for each signaling pathway between PD and CN, and found that some pathways like PSAP, VTN and SEMA4 pathways were turned off in PD, while the NRXN, NEGR, CNTN, NGL, EPHB, AGRN and CXCL pathways were turned on only in PD (Fig. 5c). Moreover, we studied the detailed changes in the outgoing and incoming signaling across all pathways using pattern recognition analysis. Four pathways were specifically active in PD, including known nerve cell adhesion signals NRXN, NEGR, CNTN and NGL, suggesting that these pathways might critically contribute to disease progression. We also found that all PD turned on pathways maintaining outgoing and incoming patterns in ENDO_Rbfox1^High^ cells. In addition, four significant pathways (NRXN, NEGR, CNTN and NGL) exhibited the most prominent outgoing and incoming signaling patterns in AST1_Meg3^High^, AST_Luzp2^High^ and AST_Lsamp^High^ cells (Fig. 5d). Corresponding to the signaling pathway, we also identified the PD up-regulated LR pair NRXN3-NLGN1, NRXN1-NLGN1 participating NRXN pathway and Negr1-Negr1 in NEGR pathway, contributing to the communication among almost all AST1 and ENDO subclusters, especially autocrine and paracrine between AST_Meg3^High^ and ENDO_Rbfox1^High^ cells (Fig. S9)

**Fig. 5.**
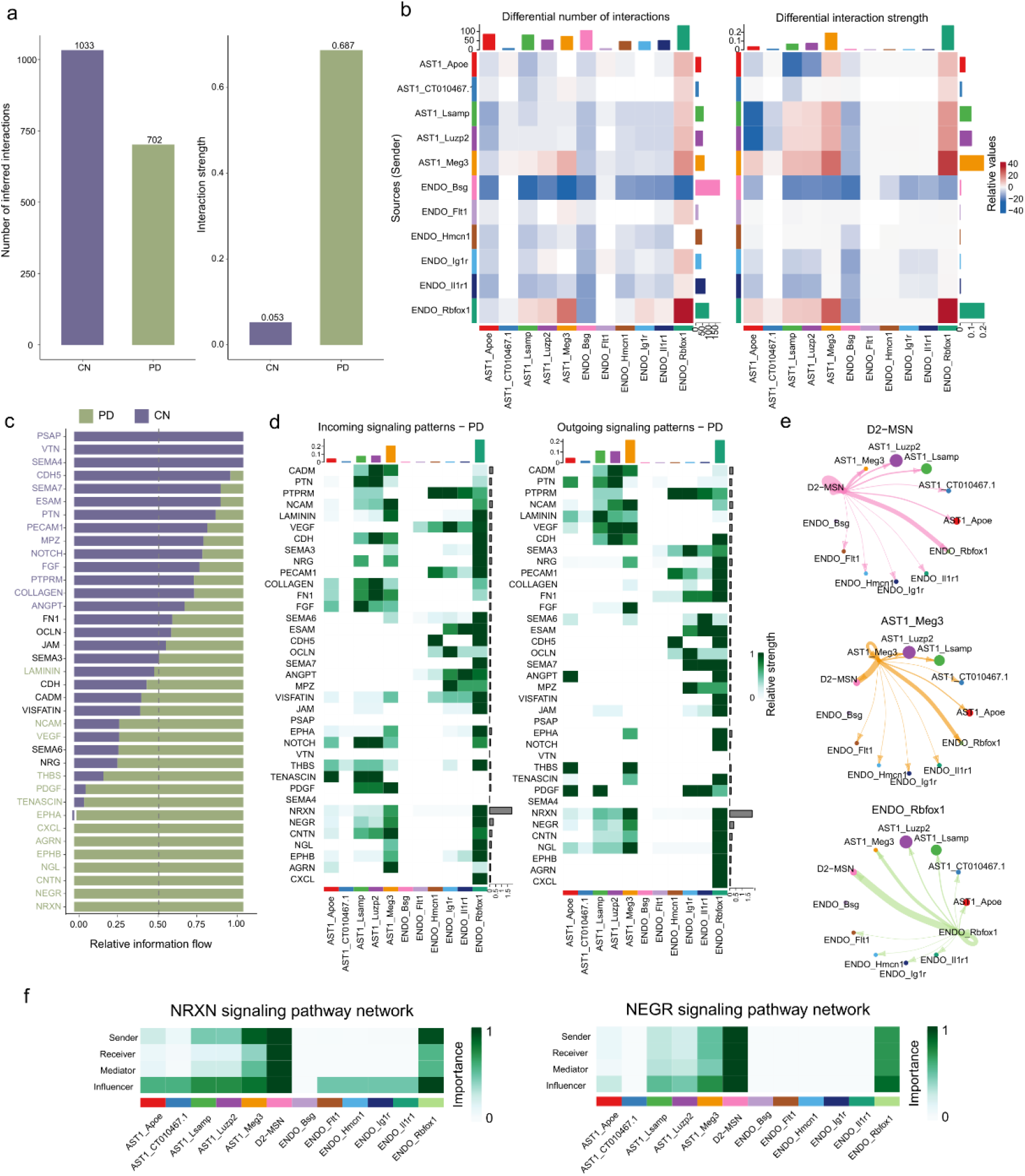
Characterization of cell communications among PD-specific cells. **a,** The number and strength of AST1 and ENDO intercellular ligand-receptor interactions in PD and CN. **b,** Heatmaps of the interaction quantity (left) and strength (right) between AST1 and ENDO subpopulations in PD and CN. **c**, Identification and visualization of conserved and specific signaling pathways. **d**, Heatmaps of the outgoing and incoming signaling patterns of AST1 and ENDO subclusters in PD. **e**, Circle plots show and compare cell–cell communication alterations among PD-specific cells. **f**, Heatmaps of the NRXN and NEGR signaling network displaying relative importance of each cell group ranked.

In order to explore the communication network among D2-MSN, AST1 and ENDO subclusters, we conducted cell communication analysis on PD data alone. We found that D2-MSN communicated with almost all AST and ENDO subclusters, especially with AST_Meg3^High^ and ENDO_Rbfox1^High^ (Fig. 5e, Fig. S10a and Fig. S11). In addition, almost all AST subclusters contributed more receptors for D2-MSN, especially AST1_Meg3^High^, AST1_Lsamp^High^ and AST1_Luzp2^High^ cells (Fig. 5e, Fig. S11). The outgoing and incoming signals contributed equally and strongly between D2-MSN and ENDO_Rbfox1^High^, indicating that these two cell types were more closely interlinked (Fig. 5e, Fig. S11). In-depth exploration of the NRXN and NEGR signaling pathway indicated that these two signaling factors play a key role in the communication network between D2-MSN, ENDO_Rbfox1^High^, and AST1_Meg^High^. D2-MSN and ENDO_Rbfox1^High^ exhibited high expression of the sender, receiver, mediator, and influencer, while almost all AST1 cells acting as an influencer (Fig. 5g, Fig. S10d). Notably, we observed that the communication probabilities of NRXN3-NLGN1, NEGR1-NEGR1 and CNTN1-NRCAM interaction were more significant (p-value < 0.01) between these three cells (D2-MSN, ENDO_Rbfox1^High^ and AST_Meg3^High^) (Fig. S10c), suggesting that MPTP induced huge cell communication network changes between these three PD-risk clusters by increasing NRXN3-NLGN1, NRXN1-NLGN1 and NEGR1-NEGR1 expression and enhancing NRXN and NEGR signaling pathway. Our analysis suggested that the alteration in intercellular communications involving AST, ENDO and MSN cells might be a previously underestimated aspect of PD pathogeneses, providing a basis for further exploration of PD mechanisms.

## Discussion

In this study, we performed snRNA-seq combined with advanced bioinformatics analysis to explore the effects of MPTP on cell states in the mouse brain. We found PD-related cell population alterations, including AST1 and ENDO, and observed changes of their activation states. We also excavated some candidate TFs and genes that might be disease-related. In addition, we identified a PD specific D2-MSN with significant changes in cellular status and gene expression. Finally, we observed enhanced cell-to-cell communication of these cells in PD. Our analysis of MPTP-induced PD mouse brain provides a reference for understanding the cellular heterogeneity underlying disease pathogenesis.

MPTP has been shown to cause pooling of blood in the brain microvasculature and decrease the permeability of the BBB, and BBB dysfunction is involved in the course of PD. BBB is mainly composed of ENDO, pericytes and AST, and reactive gliosis is a common feature of AST during BBB destruction^60^. In our dataset, we identified PD-specific astrogliosis and found its activation status and activation of TFs in disease. We observed that the AST1 subpopulations transition from a resting to an activated state with the transition from healthy to disease. It has been reported that AST is characterized by a stellate shape with multiple processes and ramifications, and become activated following brain injuries and degenerative diseases^12^. In addition, we found some potential PD marker genes associated with AST that are linked with cognitive impairment, and the highly expressed genes (Luzp2 and Lsamp) of the activated subclusters in PD were distributed in the hippocampus in Allen Brain Atlas. Although it has been demonstrated that AST can be activated in the striatum of PD^12^, hippocampus is also implicated in the cognitive dysfunction seen in some patients with PD. The lack of similar immunofluorescence experiments for verification is a deficiency of this paper, but we boldly speculated that PD-related AST cells might from or part of the hippocampus. Finally, we found that AST1 activated cell subpopulations in PD associated with endocannabinoid trans-synaptic and glutamate receptor signaling pathway. The endocannabinoid system can modify dopamine transmission trough glutamatergic synapses, and its signaling pathway is involved in the pathophysiological process of MPTP-inducted PD^61^. Evidence suggests that glutamate excitotoxicity may also play roles in the neurotoxicity of MPTP^62^. We also found heat-shock proteins (HSPs, Hspa1a) overexpressed in AST1 of PD, which is consistent with PD-specific microglia in PD human midbrain^8^. HSPs have been shown to be protective toward each of the hypothesized mechanisms of MPTP toxicity^62^. HSPs, as specific molecules produced by AST, may be a promising neuroprotective strategy in neuropathology. Therefore, we speculate that our AST1-related genes will provide important clues for PD research.

TFs tightly control cell fate in neurodevelopment and have been implicated in neurodegenerative processes. We found that most of the PD-specific activated TFs were distributed in AST1, and then identified the TFs most linked AST1 in the disease model: Dbx2 and Sox13. Studies have shown that Dbx2 encodes developing brain homeobox protein 2, highly expressed during neuronal development and regulating differentiation of interneurons in brain and spinal cord^63^. The widespread Dbx2 expression can effect on gross motoric function in fruit fly and mice^64^. In addition, Dbx2 has recently been shown to act as a TF regulating the maturation of cultured AST^65^. Similarly, the Sox gene family function as important transcriptional regulators of glial development in the central nervous system^66^. Subsequently, we clarified that these TFs play inhibitory or rewarding roles in disease, which had not been reported in any previous studies, and our analysis provides potential target markers for the treatment of PD.

The BBB is characterized by the presence of tight junctions between ENDO cells, and the expression of specific polarized transport systems, and some studies have shown alterations in ENDO tight junctions during PD development^12^. Our multichannel analysis showed that MPTP-induced PD mice were closely related to ENDO cells. We simulated the activation trajectory of ENDO cells and detected the specific deletions of reactive ENDO in PD, especially in the subset of cells of Bsg^High^. Bsg plays a crucial role in angiogenesis, and its co-expressed genes tended to be enriched in gene terms of the extracellular matrix, cell adhesion, and cell-cell interactions^67^. We hypothesized that the loss of ENDO cells in PD resulted in the destruction of tight junctions between cells, thus enhancing the permeability of BBB and promoting the entry of MPP^+^ toxins into brain environment. In addition, we also explored some genes in ENDO cells that affect neurological diseases and BBB, which might be used as candidate markers for PD diagnosis. Among them, Cxcl12 level may be potential biomarkers of inflammation in PD patients, and Slco1a5, Cldn5 and Ly6a genes are all associated with the BBB^68^. Furthermore, we found low expression of HSPs in PD-specific ENDO cells, contrary to AST1. Rise in HSPs level confers tolerance to energy deprivation, which is one explanation for the neurotoxic effects of MPTP, and our results also confirm this view^62^.

Our snRNA-seq data also showed a trend towards increased cells of OLG and OPC in PD, but our trajectory analysis results did not observe a significant PD risk association for OLG and OPC, which was consistent with the results of the latest genome-wide association studies^69^. This suggests that MPTP intake may not be the driving factor for the changes in OPC and OLG cell status. Namely, MPTP can induced more genes expression and cell state changes in AST1 and ENDO than that in OLG and OPC.

Although neuronal cells should have been one of the focuses of our study, in-depth analysis was not carried out due to the lack of more accurate information to reveal their identity. But we still found a PD-specific neurons and then decrypted its identity and status. We discovered a subcluster14 with high expression of Rarb, Pde7b, Rgs9, Gnal and unique to PD, which was defined as D2-MSN in the striatum. PD pathologies to the malfunction of the nigrostriatal dopamine pathway, where dopaminergic neurons release dopamine from axon terminals to the MSN in the dorsal striatum^2^. Although D2-MSN cells should also be detected in CN, our data separated D2-MSN only contain PD samples. We observed that D2-MSN linked genes were associated with morphine and cocaine addiction pathways. 4’-Methyl-alpha-pyrrolidinopropiophenone (MPPP) is related to morphine, piperidine and other drugs, while MPTP is an impurity in the production process of MPPP. The opioid system is involved in the reinforcing phenomenon induced by many drugs such as cannabinoids, cocaine, nicotine, but also alcohol. Imaging studies have shown that the opioid system is involved in pain processing, but also in addiction, neuropsychiatric manifestations, feeding and food disorders and, finally, movement disorders and levodopa-induced dyskinesias^70^. Dopaminergic neurons in the substantia nigra are well known to be selectively vulnerable to the MPP^+^ effects, so we speculated that our PD unique D2-MSN might be subjected to different levels of toxin attack and get huge change of cell state and gene expression.

Most scRNA-seq studies of PD mainly focused on the cell-type-specific gene expression patterns. To our knowledge, there has been rare studies characterizing the cell-cell communication with scRNA-seq data in PD research. At the most analytical stage of this work, we observed that injection of MPTP reduced the quantity of communication among AST, ENDO, and D2-MSN cells but increased in the intensity of interaction, which may be related to the energy conservation hypothesis of the PD mechanism^62^. The results of CellChat analysis not only further confirms the accuracy of our identification of PD-specific cell types, but also revealed the synergistic communication relationship between these cells. We found signaling pathways of PD-specific cells, including NRXN, NEGR, CNTN and NGL, and their roles in PD have not been reported in literatures. NRXN and NLGN are trans-synaptic proteins involved in vascular biology, and the synaptic proteins of the NRXN family are involved in the vascular system through their interaction with a basic vascular cell^71^. Studies have found that NRXN-NLGN linked synaptic function to cognitive disease^72^, and the mutations in the NRXN-1 and CNTN4 gene have been reported to cause autism spectrum disorders (ASD)^73^. NEGR1 is a generic risk factor for multiple human diseases, including obesity, autism, and depression^74^. It has been reported that NRXN and NLGN proteins are not suitable biomarkers for AD synapse pathology, but in our data their pathways are closely related to AST and ENDO in PD. We speculate that they may be potential markers for PD pathology and may be related to MPTP intake.

In summary, our study revealed several aspects of PD pathology caused by MPTP. Initially, we identified a disease-specific upregulation of AST as well as loss of ENDO cells, and systematically catalogued candidate target genes and TFs that might be associated with PD. In addition, we discovered a D2-MSN cell that exists only in MPTP-PD, which is an independent cell state initiated during MPTP-induction. Finally, the cell-cell communication between PD-specific cells was investigated in detail, and identified the PD-related signaling factors and L-R pairs. Taken together, our work at least partially supports the changes of disease-specific cells and genes in MPTP in mouse brain, and provides a useful resource and an important reference for the mechanism explanation and diagnosis and treatment of PD.

## Methods

### Ethics statement

The study was approved by the animal ethical and welfare committee of Zhongda Hospital Southeast University. All procedures were conducted following the guidelines of the animal ethical and welfare committee of SEU. All applicable institutional and/or national guidelines for the care and use of animals were followed.

### Tissue dissection and nuclear extraction

Eight-week-old male MPTP-induced Parkinson model mice (on a C57BL/6J background, MPTP-PD) and recommended control (C57BL/6J, CN) were purchased from the Shanghai Model Organisms Center, Inc. The animals were anesthetized with 500 mg/kg tribromoethanol (Sigma, Saint Louis, USA) and were killed by cervical dislocation. After the animals were sacrificed, brain tissues (cerebral cortex, hippocampus, striatum and cerebellum) were isolated, quickly frozen in liquid nitrogen and stored in liquid nitrogen until library construction. Nuclei were isolated by adapting the published 10X Genomics^®^ protocol for ‘Isolation of Nuclei for Single Cell RNA Sequencing’. In brief, the tissue was lysed in a chilled lysis buffer (10 mM Tris-HCl, 10 mM NaCl, 3 mM MgCl2, 0,1% NP-40). Then, the suspension was filtered and nuclei were pelleted by centrifugation. Nuclei pellets were then washed in ‘nuclei wash and resuspension buffer’ (1x PBS, 1% BSA, 0.2 U/μL RNase inhibitor, 2 mM DTT), filtered and pelleted again. Cell count was then performed to calculate the concentration of nuclear suspension.

### Library construction and sequencing

Sorted nuclei were processed using the 10x Chromium Next GEM Single Cell 3’ Kit v3.1 to generate the cDNA libraries. The quality of cDNA was assessed using the Agilent 2100 Bioanalyzer System. Sequencing was performed on Illumina NovaSeq 6000-S2.

### Data demultiplexing and quality control

We first used Cell Ranger 5.0.1 (10×Genomics) to process raw sequencing data, and then Seurat v4.0 was applied for downstream analysis. Before we started downstream analysis, we focused on four filtering metrics to guarantee the reliability of our data. (1) Genes detected in less than three cells were filtered to avoid cellular stochastic events; (2) Nuclei with a percentage of expressed mitochondrial genes are greater than 10% were removed to rule out apoptotic cells; (3) Cells with UMI greater than 10,000 were removed to filter out the doublet-like cells; (4) Cells with detected genes out of the range of 200-4,000 were removed. After filtering cells and genes according to the metrics mentioned above, we further applied Doublet Finder V2.0 with default parameters to predict and remove potential doublets within each sample. All in all, there are 22431 genes and 19531 cells left for downstream analysis.

### Clustering and cell annotation

After quality control, unsupervised clustering was performed using Seurat v3^75^ in a region-independent fashion. A series of pre-processing procedures including normalization, variance stabilization and scaling data, were performed in an R function ‘SCTransform’ based on regularized negative binomial regression. Then, we selected 3000 highly variable genes were selected to integrate all sequencing libraries (including PD and CN) using ‘FindIntegrationAnchors’ and ‘IntegrateData’ functions, followed by the regression of technical noise. Principal component analysis (PCA) was performed using integrated output matrix, and principal component (PC) significance was calculated using the ‘JackStraw’ function. In this case, we chose the top 30 significant PCs for downstream cluster identification and visualization. Clusters were defined based on ‘FindClusters’ function with resolution equals to 0.8. After the primary clustering analysis, we found a high proportion of excitatory neuron having high gene expression similarity. Therefore, we applied 2-rotation cluster strategy. Briefly, After the first clustering analysis, we obtained major cell types, then we sub-clustered the excitatory neurons with resolution equals to 0.25 and merged the 2-rotation results as final cluster results. Uniform Manifold Approximation and Projection (UMAP) was used for the final dimension reduction and visualization.

Based on the cluster results, we next used ‘FindAllMarkers’ function with MAST algorithm which was specially developed and applied to single cell data detecting differential expressed genes to identify marker genes for each cluster. We rank the marker genes according to the p-value and log2 fold change (log2FC) within each cluster and searched top genes in Cell Marker^76^ and Panglao DB^77^ databases to annotate cell types of clusters.

### Differential expressed genes analysis and gene set functional annotation

Within each cluster, we detected differential expressed genes (DEGs) between PD and CN conditions by using ‘FindMarkers’ function. Use setted to ‘MAST’ as well and controlled false-discovery rates (FDRs) using the Benjamini-Hochberg procedure. Then we set threshold q_adjust < 0.05, abs | log2FC | > 1 to filter DEGs and got PD- up and -down regulated genes compared to CN for each cluster.

The DEGs functional enrichment analysis based on GO and KEGG was applied by an R package ClusterProfile^78^ v3.18.4 which using a hypergeometric test and corrected for multiple hypothesis by FDR.

We used R packaged wordcloud2 to show the frequency of PD-risk genes detected in differentially expressed genes within each cluster, the bigger word size indicates the more frequency.

### Inference of regulon, quantify cell-type specificity score and functional validation

To predict gene regulatory networks using single nuclei gene expression data, we used pySCENIC^18^ approach. There are three major steps of SCENIC to construct high confident gene regulatory networks. First of all, SCENIC calculate co-expression modules between TF and candidate target genes using GENIE3. Then RcisTarget was used to create regulon with only direct targets by identifying modules for which the regulator’s binding motif is significantly enriched across the target genes. Finally, AUCell scores each regulons active score in each cell and create a binarized activity matrix between regulons and cells. Using this matrix, we can predict cell states without removing batches and identify cell type specific activate regulons.

For identifying cell state specific regulons, we adapted an entropy-based method to quantify cell-type specific score of each regulon. Firstly, we name the vector of 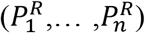 as P^R^ to describe the distribution of regulon activity score, the vector of 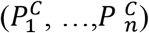 as P^C^ which can indicate whether a cell belongs to a specific cluster, n is the total cell numbers. Then we calculate the Jesen-Sannon Divergence (JSD) from P^R^ and P^C^:

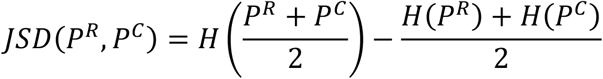

where*H*(*P*) = −Σ*p_i_loa*(*p_i_*)

Finally, the regulon specificity score (RSS) is calculated as this:

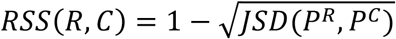

Therefore, we know the range of RSS is (0,1), and if the regulon activity is highly different among clusters the RSS approach to 1, otherwise if there is no difference of regulon activity among clusters the RSS will be equal to 0.

To further validate whether our predicted regulons are functional related to their associated cell types or PD condition specific activated, we employed an online tool SEEK^79^. SEEK provides the gene co-expression search function for lots of mouse database from the GEO (Gene Expression Omnibus), so we can detect that whether the genes within the same regulon are co-expressed and which kinds of papers data had also reported the similar co-expression module. If genes within regulon are significantly co-expressed in many datasets related to Parkinson disease or some certain cell types, it could as a further evident that the regulon was reported to be highly related to the Parkinson’s disease in certain cell type.

### Trajectory analysis using Monocle3

To obtain cellular state changes between PD and CN samples within AST1, ENDO, OLG, OPC and part of the excitatory neurons, we reconstructed the cellular states trajectories using the standard Monocle3^26^ workflow. Firstly, we subdivided certain clusters, used the filtered raw counts as input to integrate PD and CN cells and normalized factor size. The sample effect was removed using the Mutual Nearest Neighbor method with parameter ‘alignment_k = 20’. The reduce_dimension function was used for dimensionality reduction, and the Louvain method was used for clustering with a resolution of 0.01. Then, the trajectory inference used the learn_graph function with default parameters. Finally, Pseudotime ordering was done by rooting the trajectory manually based on the shape of trajectory and background knowledge.

The most important step is to identify trajectory-dependent genes which may influence the PD and CN cell states slightly changes within each cluster. We first calculated subcluster marker genes using ‘topmarkers’ function then using ‘graphtest’ function which using the spatial correlation analysis Moran’s I approach to identify highly variable genes associated with the trajectory. Thus, the trajectory-dependent genes were defined by intersection of subcluster markers and trajectory associated highly variable genes.

### Cell-cell communication analysis

To further investigate the intercellular communication changes induced by MPTP, we used R software CellChat^80^ v1.4.0 to calculate communication networks between subclusters of AST1, ENDO and D2-MSN. We predicted the communication network including signaling pathway and ligand-receptor (L-R) pairs information in PD and CN samples separately and then compared the network difference between these conditions. The interaction number and strength are two key factors, so we used ‘compareInteractions’ function to get whole network interaction number and strength differences. Then, for the conserved signaling pathways, we ranked these pathways according to their Euclidean distance in the shared two dimensions space. The top pathways indicated more difference between PD and CN. We also compared each signaling pathway’s information flow, which is the sum of communication probability among all cell pairs, to identify different pathway states including turn off/on, decrease and increase in one condition compared to the other. Finally, we zoomed in the LR pairs level, and calculate dysfunctional L-R pairs by using differential expression analysis with ‘identifyOverExpressedGenes’ and ‘netMappingDEG’ functions. The up-regulated and down-regulated LR pairs can be detected. All the plot functions are from the CellChat package.

---

**Fig. S1.**
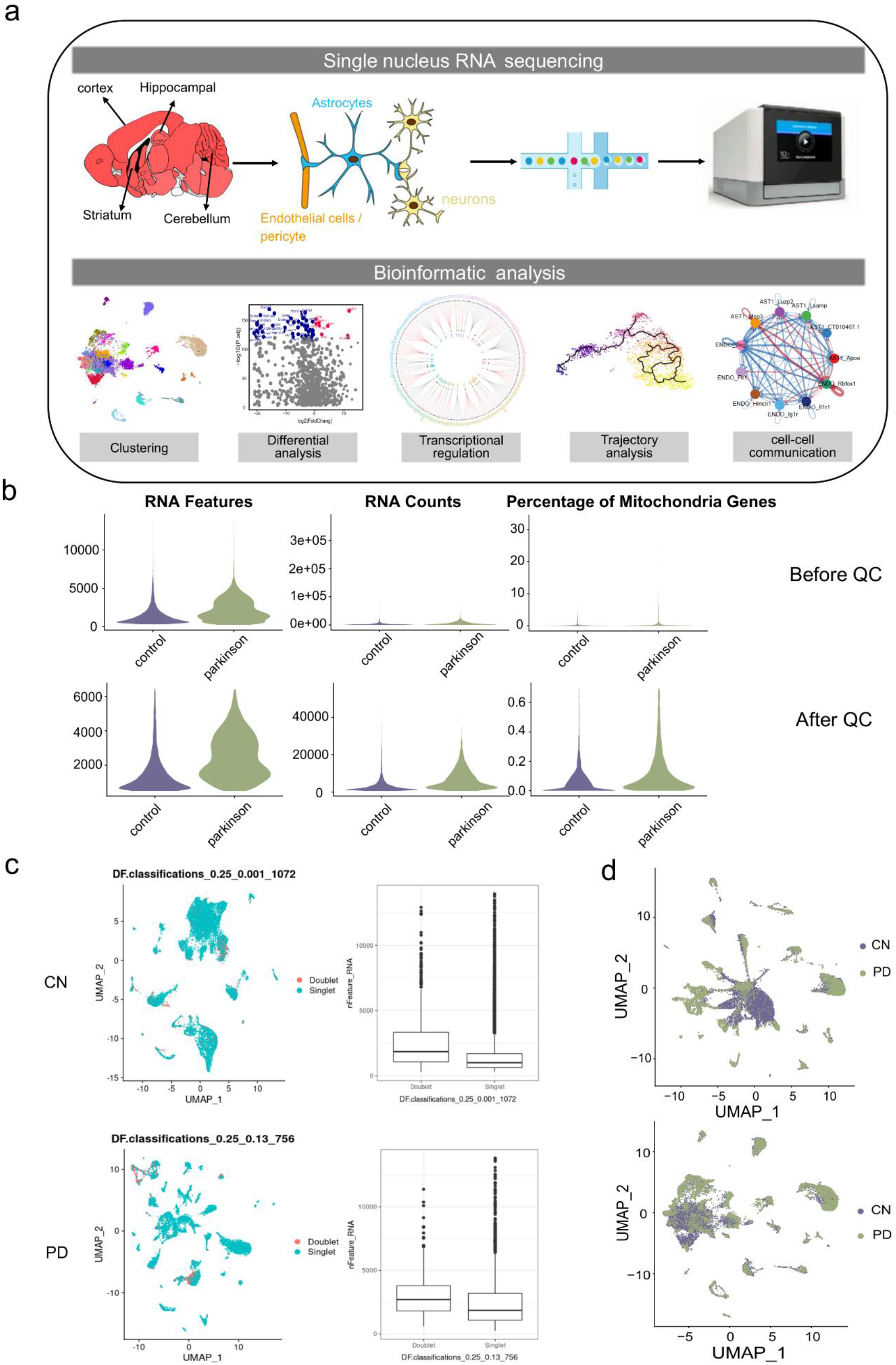
snRNA-seq quality control metrics and similarity. **a,** Schematic representation of the samples used in this study, sequencing experiments and downstream bioinformatic analyses. **b,** Information of samples before (top) and after (bottom) quality control. **c,** Potential doublets were predicted and removed by the Doublet Finder V2.0. **d,** 2D UMAP embedding of single nuclei RNA profiles before (top) and after (bottom) removing batch effect.

**Fig. S2.**
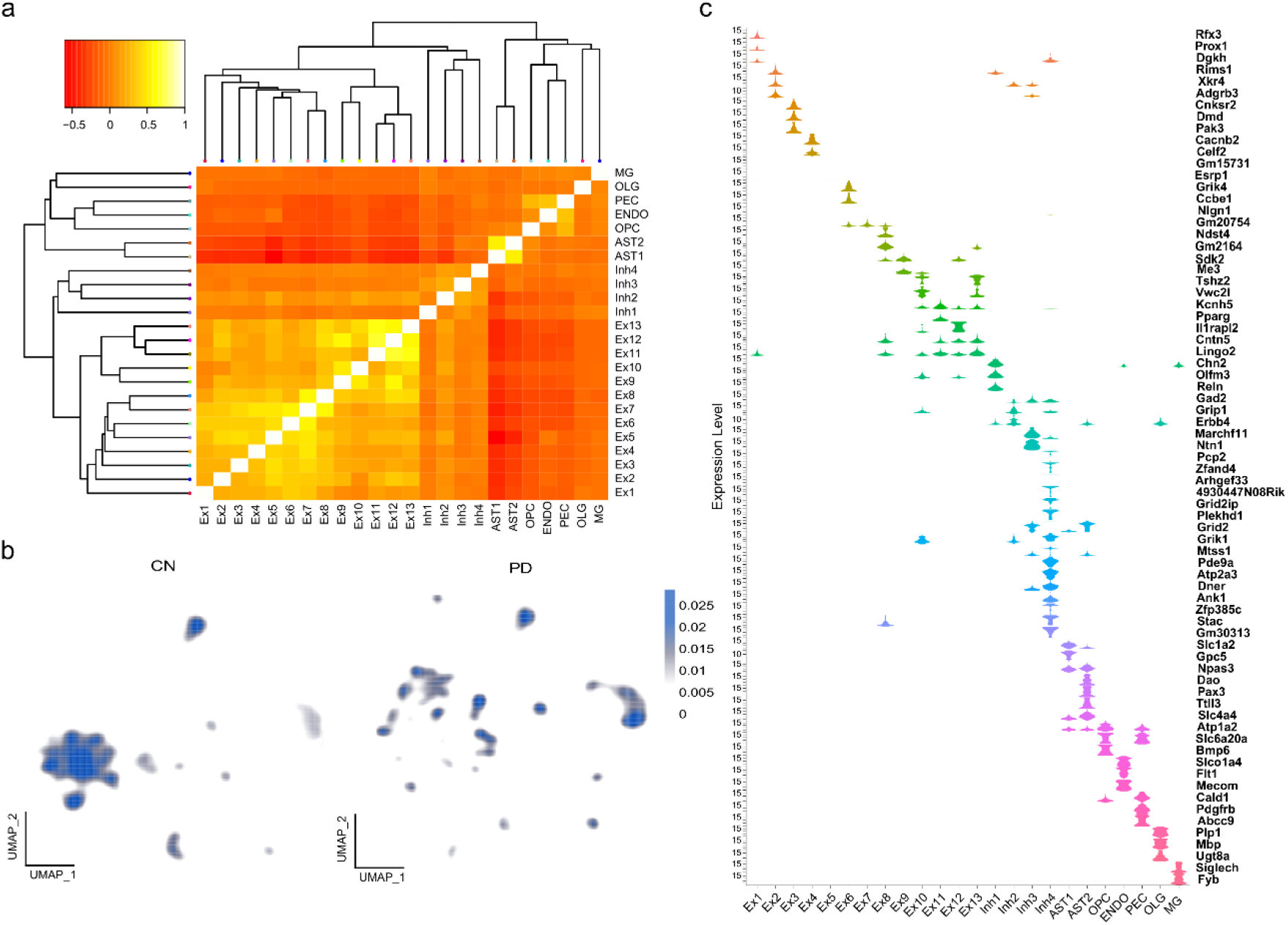
Cell type marker genes and assignments. **a,** Confusion matrix results of the machine learning cross-validation approach to validate the cell type definition. **b,** Expression distribution of cell-type marker genes. **c,** 2D cell density UMAP embeddings for PD (right) and CN (left).

**Fig. S3.**
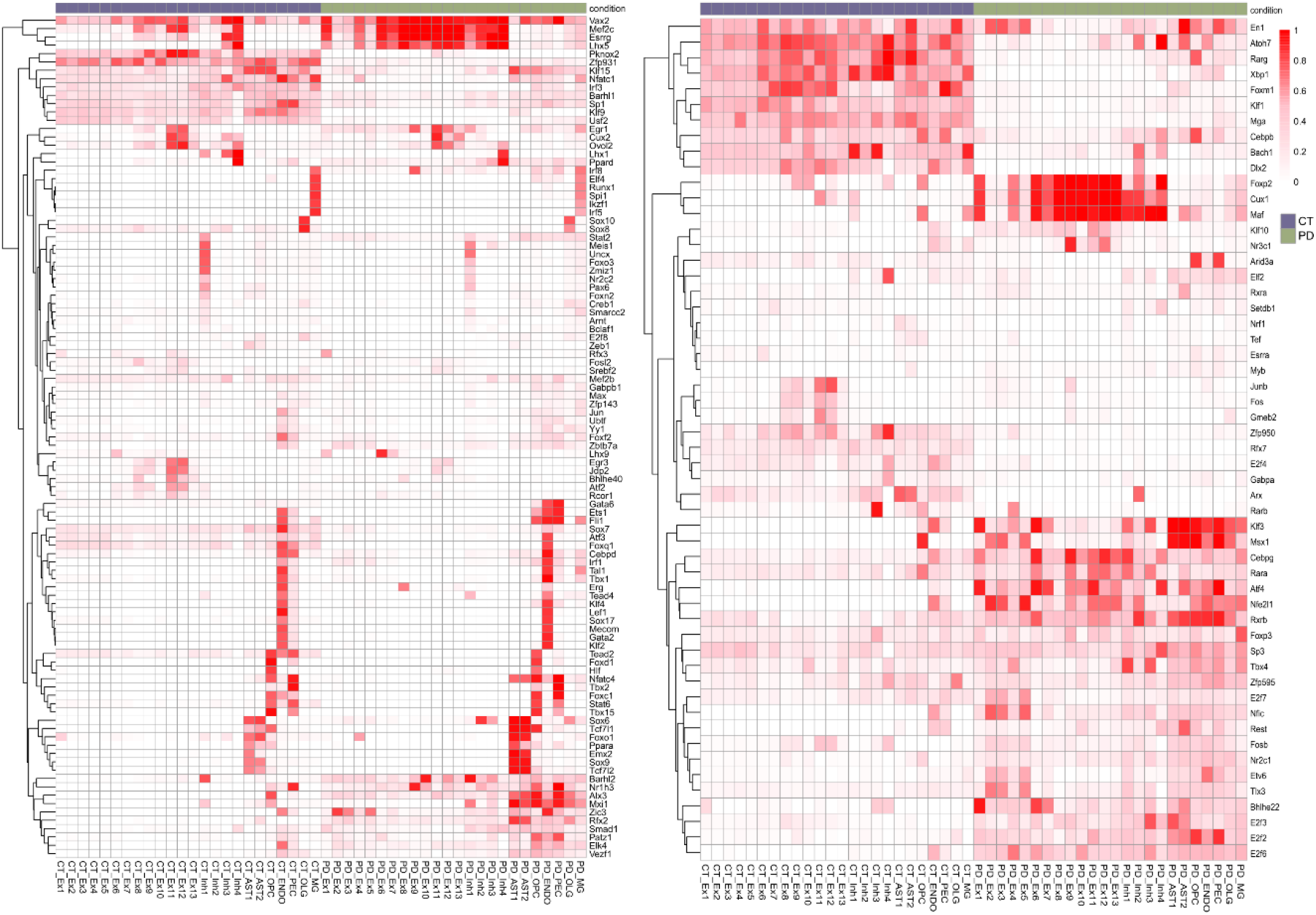
Heatmap of 155 TFs co-activated by PD and CN. There were 101 conservative TFs with similar activation status in PD and CN (left), and 54 TFs with specificity (right).

**Fig. S4.**
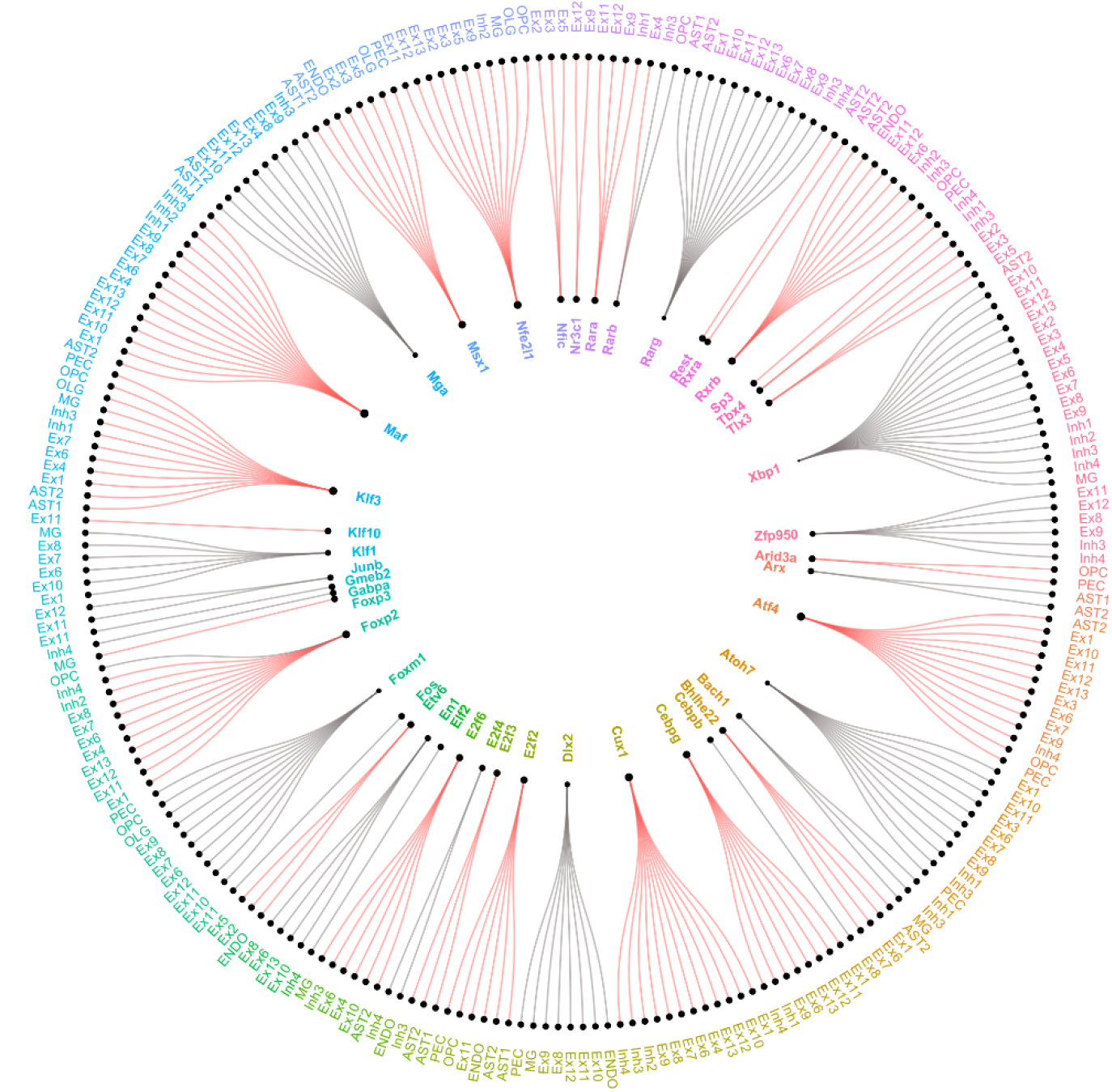
Cirle plot of 54 differentially activated TFs and their corresponding cell types in PD and CN (abs | pct (PD) – pct (CN) | >= 0.5). The inner ring is TFs, and the outer ring is a qualified cell types; The red and grey lines indicate activation in PD and CN respectively, and the size of the inner ring points indicates the number of differences.

**Fig. S5.**
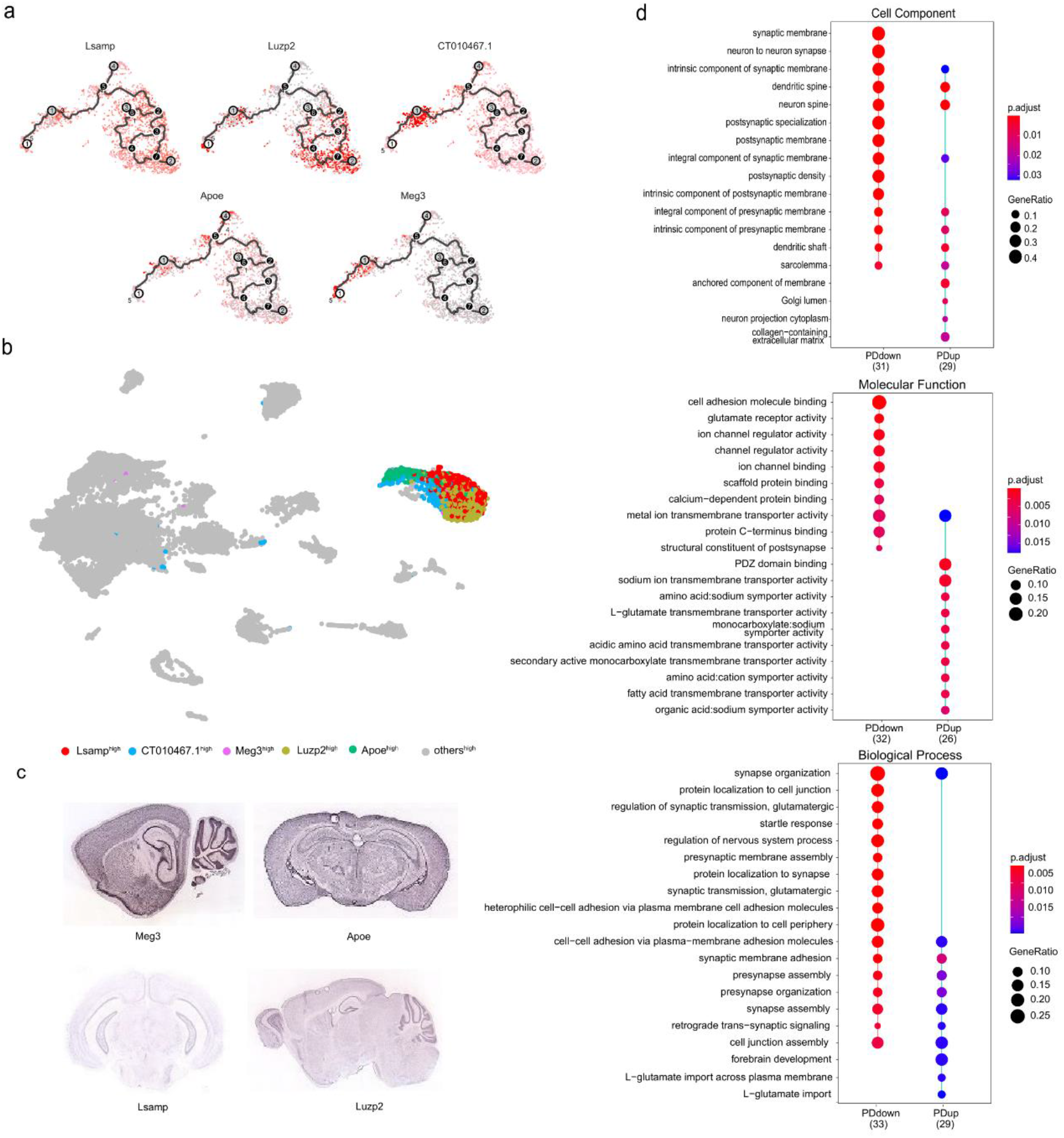
**a,** The expression of marker genes along the AST1 subclusters. **b,** UMAP embedding of five AST1 subclusters. **c,** Distribution of AST1 subclusters marker genes in Allen Brain Atlas. **d,** GO analysis of all AST1 subcluster DEGs. The color of the point corresponds to the value of p_adjust, and the size represents the proportion of the number of differential genes in the total number of differential genes under the GO term.

**Fig. S6.**
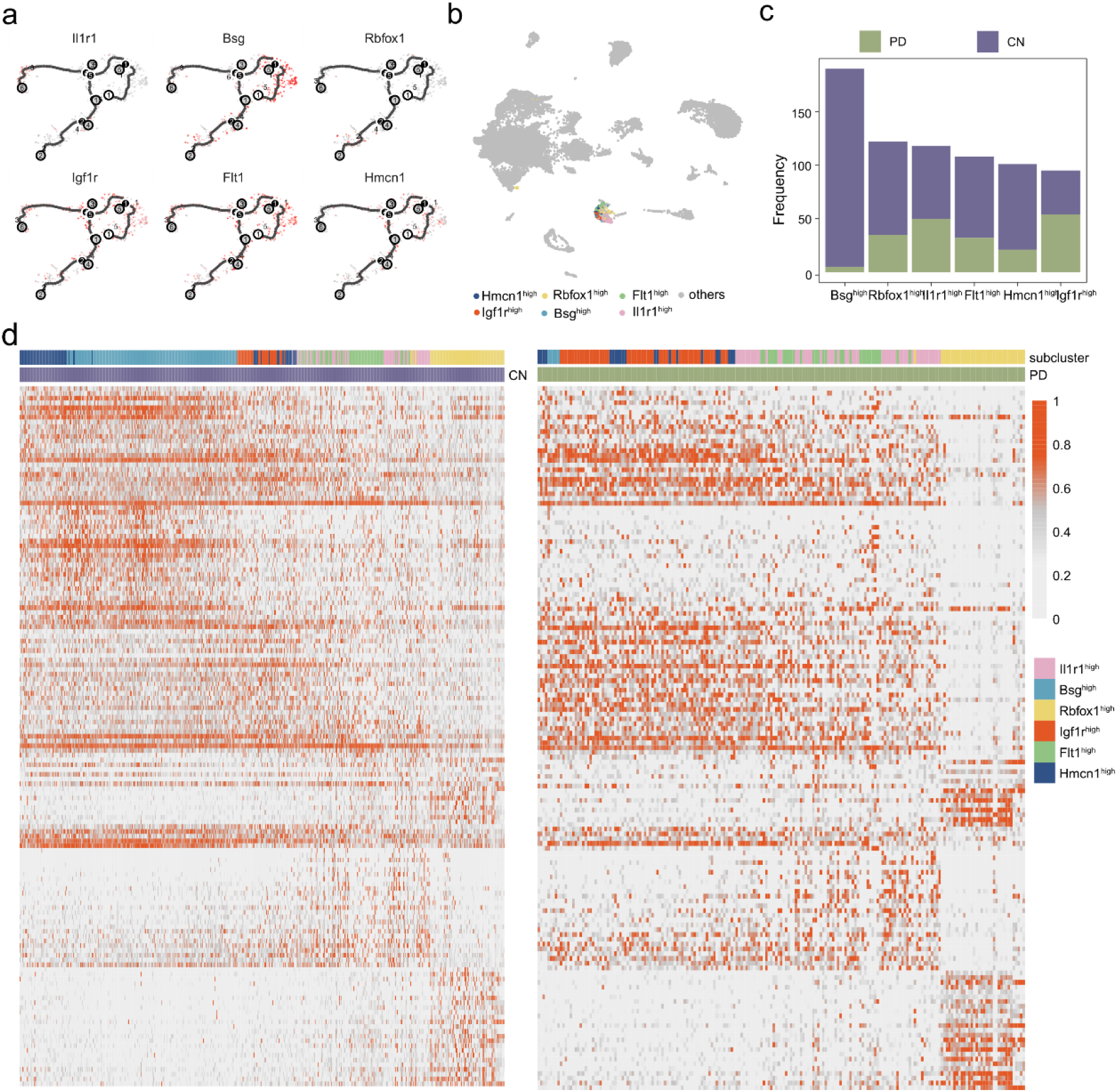
**a,** The expression of marker genes along the ENDO subclusters. **b,** UMAP embedding of ENDO subclusters. **c,** the proportion of ENDO subclusters in PD and CN. **d,** Trajectory dependent gene of each cell subcluster gene in PD (right) and CN (left).

**Fig. S7.**
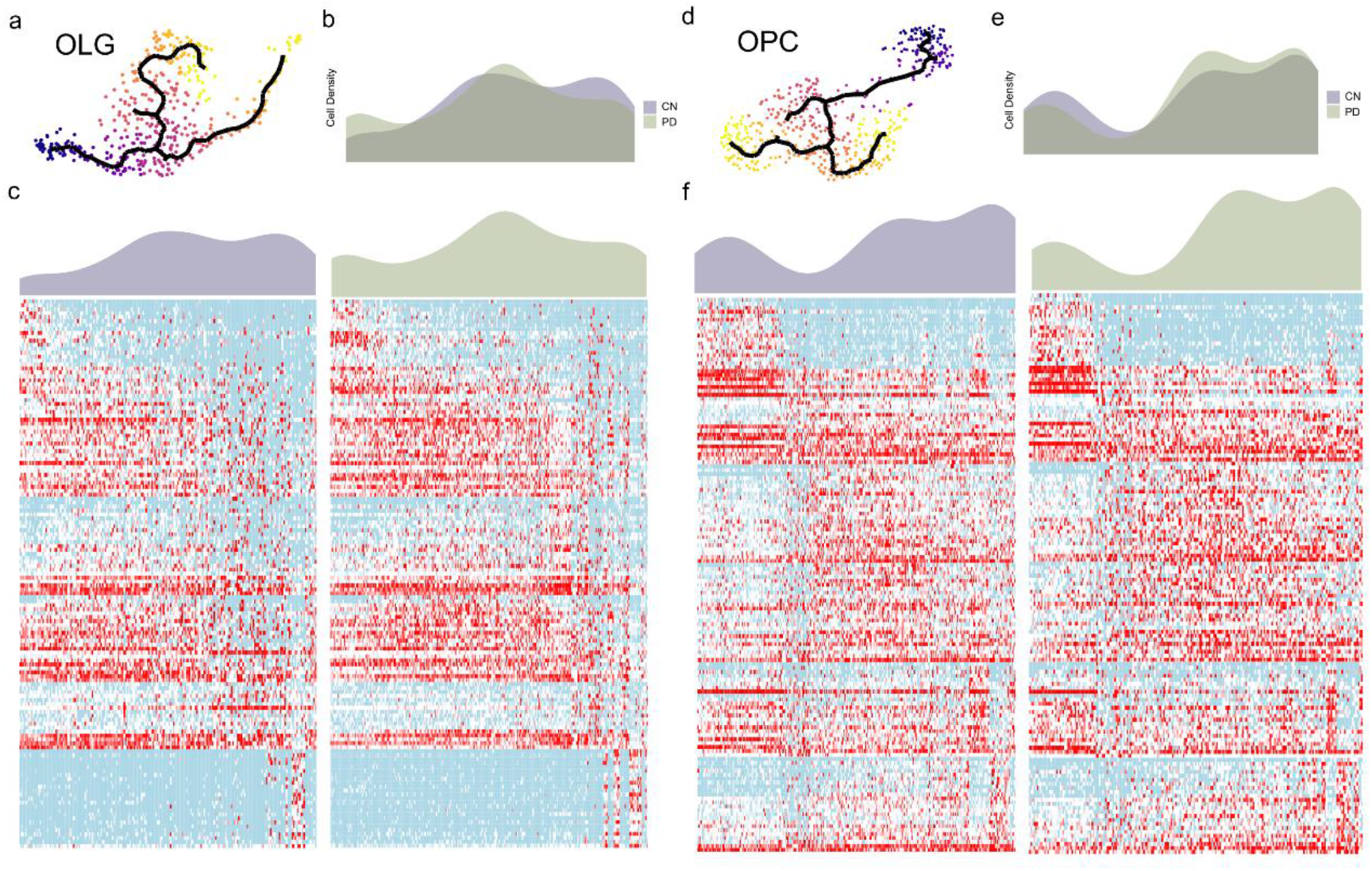
Trajectory reconstruction reveals oligodendrocytes and oligodendrocyte precursor cells differential activation in PD. **a, d,** OLG and OPC trajectory reconstruction and pseudotime representation of subclusters. **b, e,** differential cell-density distribution over pseudotime in CN and PD, separately. **c, f,** Trajectory dependent gene of each cell subcluster gene in OLG (c) and OPC (f).

**Fig. S8.**
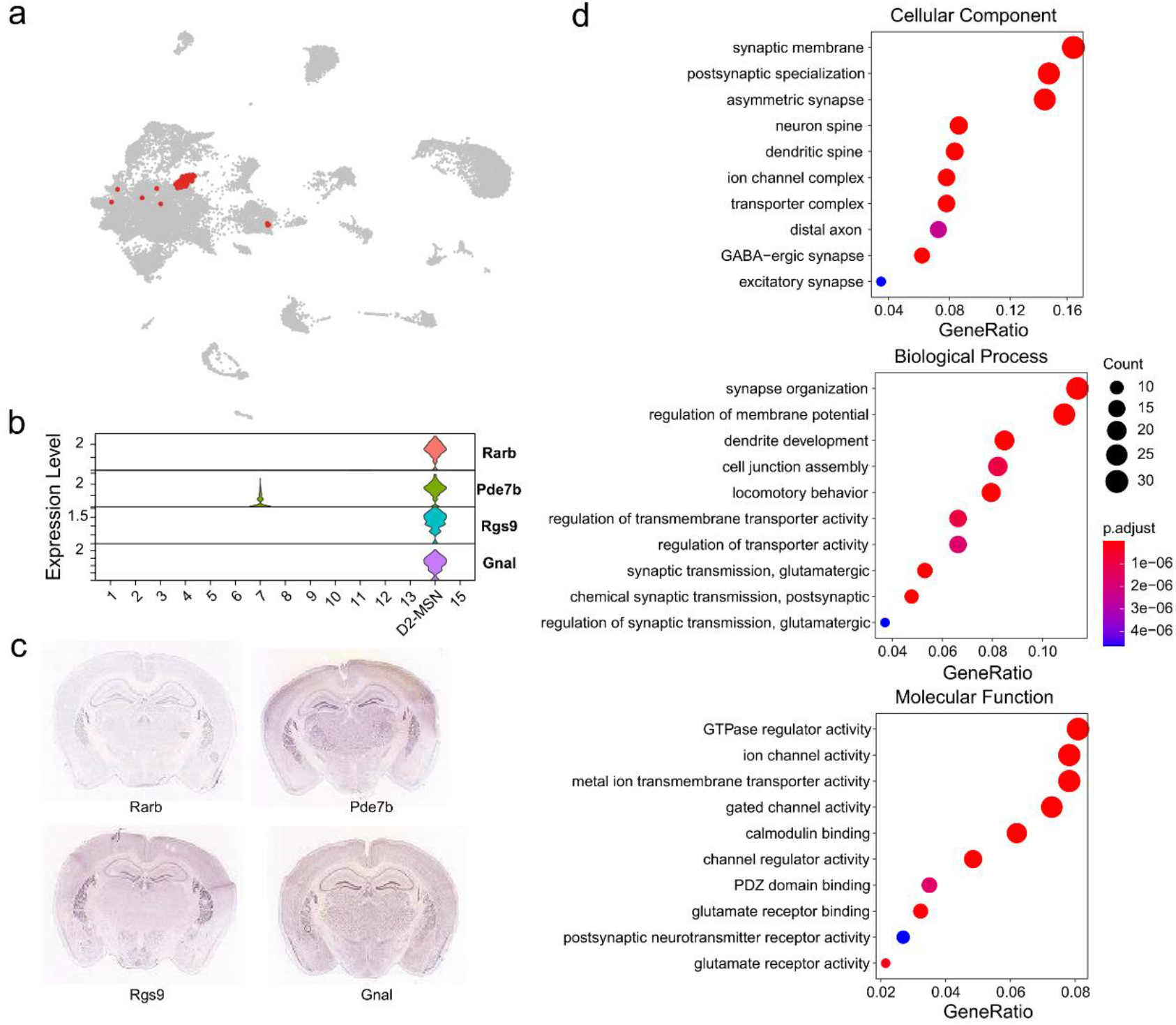
UMAP embedding of subclusters 14, which overlaps with EX4. **b, c,** The top four highly expressed marker genes of subclusters 14 in the ISH of Allen Brain Atlas. These genes are distributed in the striatum (STR). **d,** GO analysis of all marker gene expression in subcluster 14. The color of the point corresponds to the value of p_adjust, and the size represents the number of differential genes under GO terms.

**Fig. S9.**
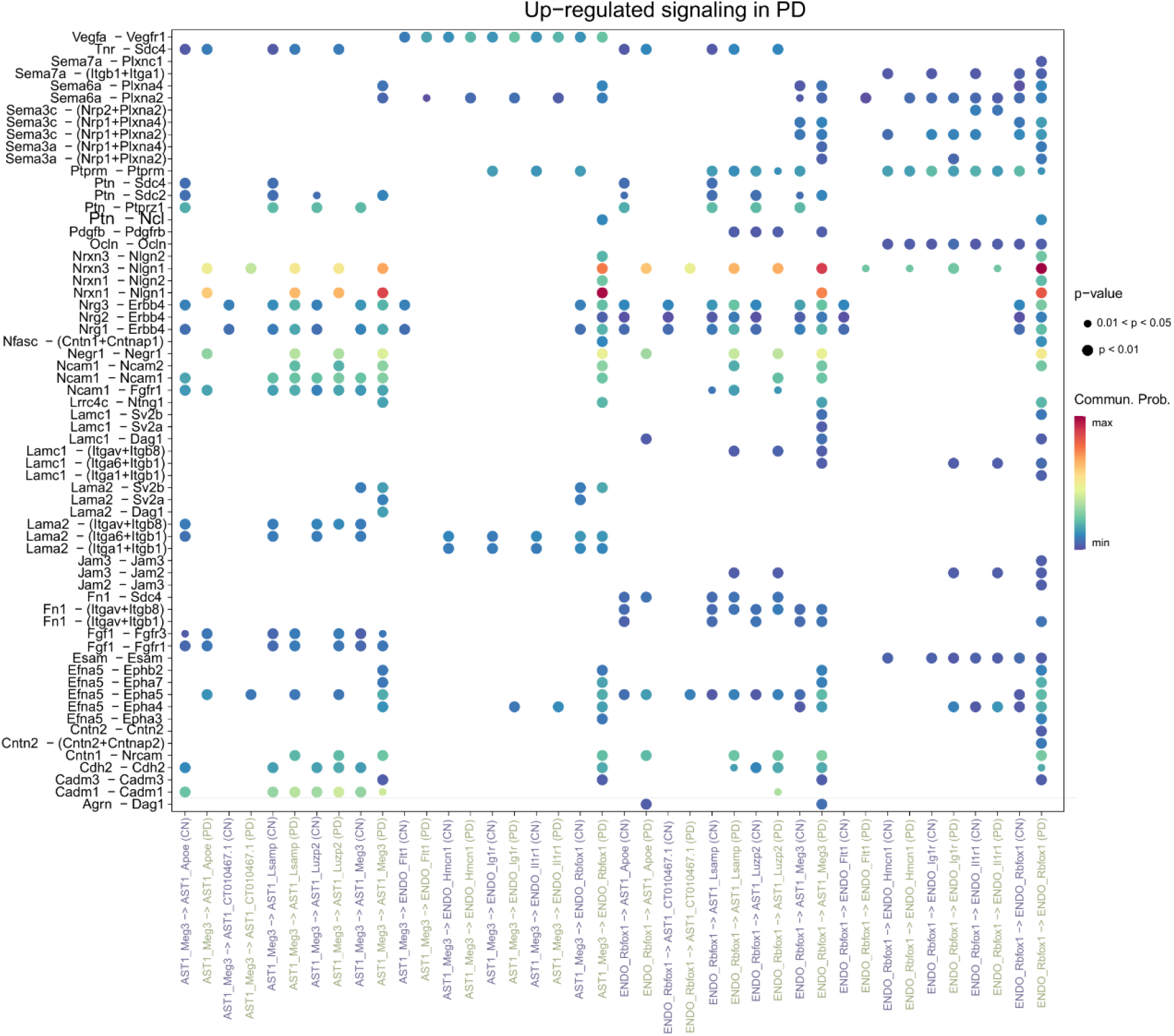
Up-regulated Ligand-receptor pairs that contribute to the signaling of PD-specific cells. The dot color and size represent the calculated communication probability and p-values.

**Fig. S10.**
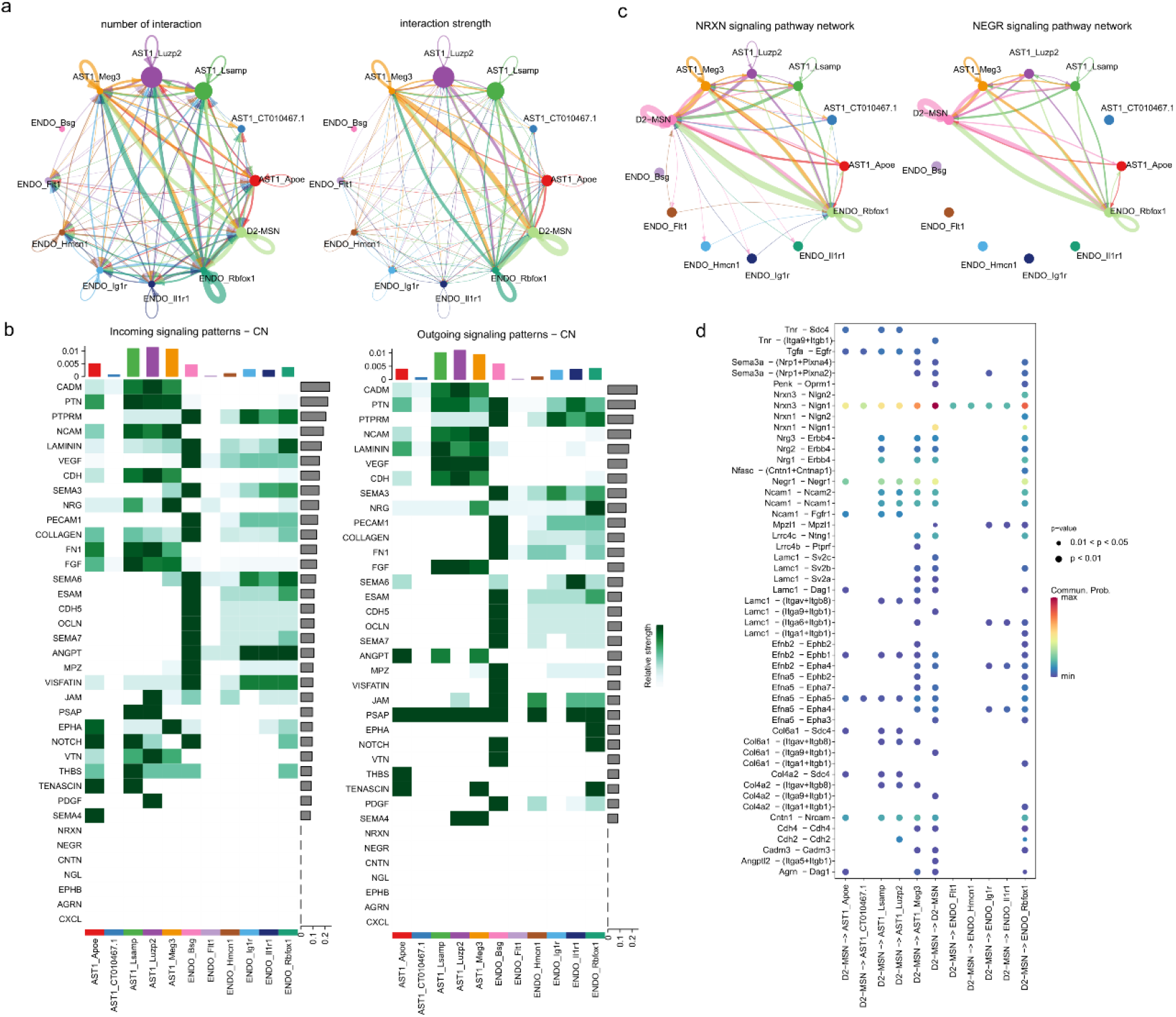
**a,** Circle plot of the number and interaction strength of ligand-receptor pairs between PD-specific cells. **b**, Heatmaps of the outgoing and incoming signaling patterns of AST1 and ENDO subclusters in CN. **c**, Comparison of the significant ligand-receptor pairs between PD-specific cells, which contribute to the signaling from D2-MSN to AST1 and ENDO subpopulations. Dot color reflects communication probabilities and dot size represents computed p-values. **d**, The inferred NRXN and NEGR signaling networks. Circle sizes are proportional to the number of cells in each cell group and edge width represents the communication probability.

**Fig. S11.**
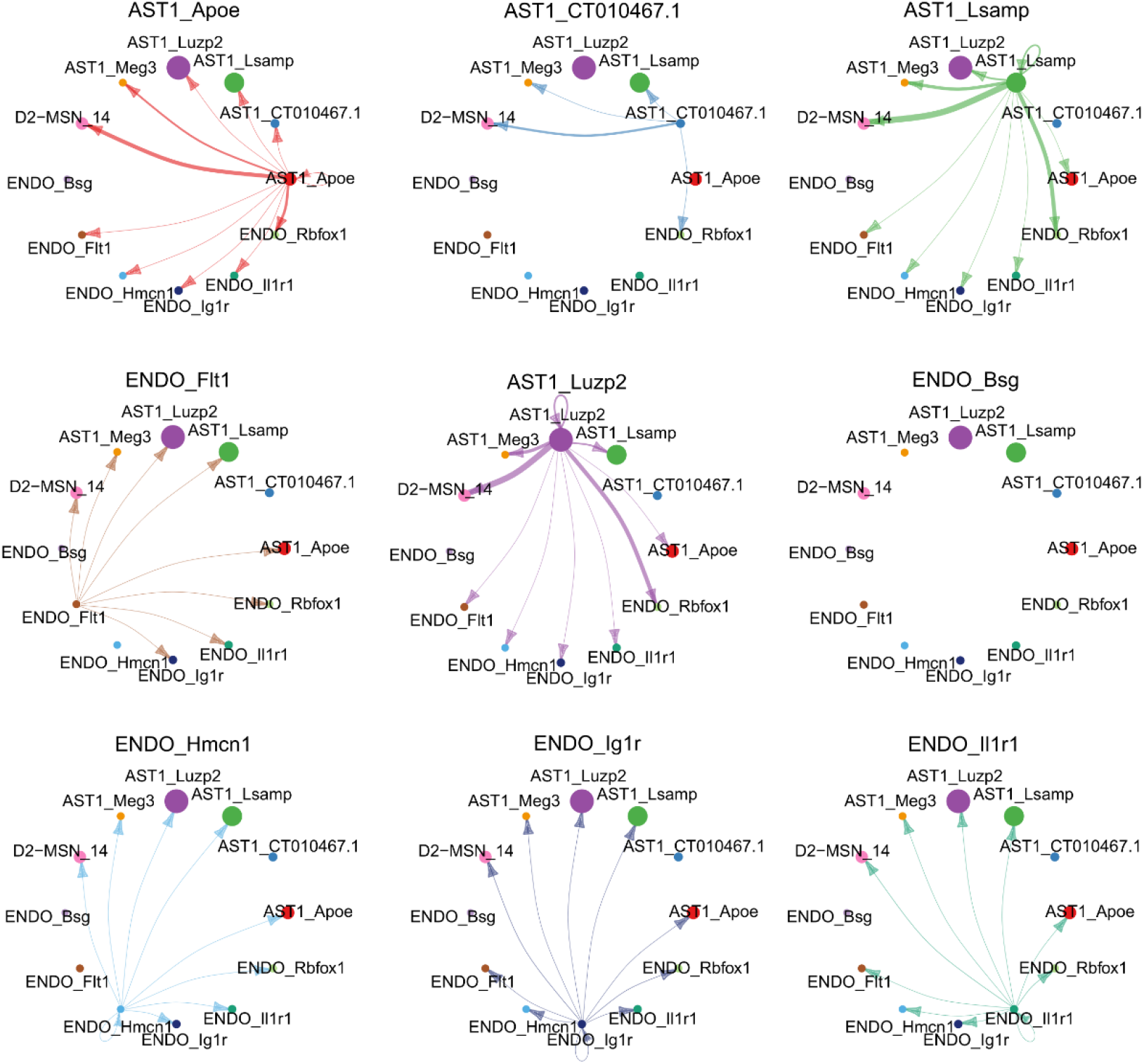
Circle plots of the interaction strength among D2-MSN, AST1 and ENDO subclusters in PD.

